# Neural signatures of associational cortex emerge in a goal-directed model of visual search

**DOI:** 10.1101/2025.06.06.658387

**Authors:** Motahareh Pourrahimi, Pouya Bashivan

## Abstract

Animals actively engage with their environment to gather information, continuously shaping both their sensory input and behavior. Understanding this closed loop between perception and action remains a central challenge in neuroscience. A key example is active vision, where observers decide where to look next, selectively sampling from their visual space to guide ongoing perception and action. However, despite major advances in linking neural activity with behavior and computational modeling of vision under passive viewing conditions, the interactive aspects of natural vision remain underexplored. Visual search, the act of locating a target among distractors, exemplifies this dynamic sampling process and has long served as a core paradigm for studying visual attention. While its behavioral and neural signatures have been characterized in humans and non-human primates, a unifying model that links these neural phenomena to behavior during visual search has been lacking. Here, we present a biologically aligned neural network model trained to perform visual search directly from natural scenes by generating sequences of saccades to locate a target. The model generalizes to novel objects and scenes, produces human-like scanpaths, and recapitulates classic behavioral biases in human visual search. Strikingly, units in the model exhibit neural response properties characteristic of the fronto-parietal network, including a stable cue template in working memory, a retinocentric cue-similarity map, and prospective fixation signals. Beyond reproducing known behavioral and neural phenomena, the model reveals a representational geometry that supports cue-driven prioritization, spatial memory, and planning of future fixations. These results establish a computational framework for studying visual search as an emergent property of goal-directed perception, offering concrete predictions for neurophysiological and behavioral testing, and paving the way toward a unified account of active vision.

## Introduction

Vision is among the most studied brain functions, yet much of our computational understanding remains grounded in passive paradigms, such as core object recognition Yamins et al. (2014); Yamins and DiCarlo (2016); Kar and DiCarlo (2020); DiCarlo et al. (2012), which capture only a narrow slice of animals’ visual capabilities. In natural environments, however, vision is not a static process—it is active, goal-driven, and tightly coupled with behavior Hayhoe and Ballard (2005); Land and Tatler (2009); Gottlieb et al. (2014). Animals continuously shape their own stream of sensory input through interactions with the environment, directing their gaze to locate food, monitor threats, or explore novel stimuli. This dynamic sampling of the world is central to how perception supports action, and it is especially crucial for survival in complex ecological settings Gottlieb and Oudeyer (2018); Yarbus (1967); Hayhoe and Ballard (2005). These behaviors are best understood under the broader umbrella of active vision, where perception is deeply entangled with the decisions that determine what is seen next. Within this framework, visual search stands out as a foundational behavior. Whether rummaging through a cluttered desk for car keys, scanning a crowded cafe for a friend, or reaching for the salt shaker at dinner, humans and other animals rely on search as a core cognitive operation Eckstein (2011); Wolfe (2021). Its pervasiveness has made it a central paradigm in psychology, neuroscience, and machine learning, both as a core cognitive paradigm and as a building block in numerous primate behaviors Treisman and Gelade (1980); Gelade (2001); Wolfe (1994); Duncan and Humphreys (1989); Eckstein (2011); Mitroff and Biggs (2014); Chan and Hayward (2013); Deco and Heinke (2007); Malcolm and Henderson (2009); Wolfe and Horowitz (2004); Egeth and Yantis (1997); Yang et al. (2017); Li et al. (2020); Yuan and Li (2020); Itti and Koch (2001); Mazer and Gallant (2003); Ischebeck et al. (2024); Desimone et al. (1995); Chelazzi et al. (1993); Wolfe and Horowitz (2017); Kristjánsson and Draschkow (2021); Bella-Ferńandez et al. (2022). Visual search is not merely about detecting targets; it exemplifies the fundamental challenge of how intelligent agents allocate limited perceptual resources to make sense of a rich, high-dimensional world.

Developing computational models that accurately capture visual search behavior and elucidate the neural computations supporting it has been a long-standing challenge. Early efforts adopted a reductionist perspective, often studying *covert* search (where eye movements are suppressed) rather than the more naturalistic *overt* search behaviors that involve active gaze control. Classical “box-and-arrow” models offered conceptual accounts of attention shifts but lacked grounding in neurophysiological detail. More recently, a range of data-driven Yang et al. (2024); Mondal et al. (2023); Yang et al. (2022); Travi et al. (2022); Yang et al. (2020); Zelinsky et al. (2019) and algorithmic Zhang et al. (2018); Rashidi et al. (2023); Ding et al. (2022); Zhang et al. (2022); Bujia et al. (2022); Gupta et al. (2021); Adeli and Zelinsky (2018); Adeli et al. (2017); Wei et al. (2016); Zelinsky et al. (2013); Pomplun and Gray (2007); Rao et al. (2002); Zelinsky (2008); Navalpakkam and Itti (2006, 2005) approaches have been developed, aiming to predict human scanpaths using machine learning or formalized hypotheses about search strategies, respectively. Data-driven models typically consist of parametrized functions that are fitted to replicate the human scanpaths from eye tracking data, without an explicit intention to reveal the underlying neural mechanisms of visual search Yang et al. (2024); Mondal et al. (2023); Yang et al. (2022); Travi et al. (2022); Yang et al. (2020); Zelinsky et al. (2019). Despite their strong performance in predicting human behavior, it remains unknown whether and how their components and learned representations correspond to brain networks involved in visual search. On the other hand, algorithmic models typically consist of rule-based or parameterized models driven by specific hypotheses about how the brain performs visual search without being grounded in biologically-plausible neural architectures. As a result, these models are unable to explicitly inform the role of different brain areas in executing visual search behavior as they lack distributed representations that can be compared with the function and response properties of neurons in the brain. Notably, none of these prior models are image-computable. Although Zhang et al. (2018); Ding et al. (2022) incorporate image-computable vision modules, they replace the neural computations beyond sensory visual areas with a symbolic rule-based algorithm. Consequently, they do not offer any concrete predictions about the neural responses in the higher-order areas such as the frontal eye fields (FEF), lateral intraparietal area (LIP), ventral prearcuate area (VPA), and the superior colliculus (SC) during visual search.

The progression of modeling core object recognition—and vision science more broadly—has followed a trajectory similar to that of visual search: beginning with descriptive box-and-arrow models, and advancing through reductionist algorithmic frameworks. Critically, it culminated in the breakthrough of convolutional neural networks that accurately predicted neural responses along the primate ventral visual stream—a milestone that the field of active vision, and visual search as an example, had long been awaiting. Since then, artificial neural networks have emerged as a powerful class of models for explaining both animal behavior and neural activity across numerous neuroscience domains, including vision, navigation, memory, learning, decision-making, and planning. These networks provide a scalable, image-computable, and flexible framework for modeling brain function and behavior, imposing fewer structural assumptions and requiring less task-specific hand-engineering than prior approaches. Recognizing this gap, we present GVSM (Goal-directed Visual Search Model), a visual search model operating on high-dimensional naturalistic visual input that is both biologically grounded and capable of performing the task from first principles.

To this end, we 1) used a network architecture capable of integrating information across fixations through a dynamically updated internal state; 2) trained the model solely on the task of performing visual search without assuming any symbolic rules and computations; 3) implemented eccentricity-dependent visual acuity similar to that in primates via eccentricity-dependent downsampling of the input to simulate the sensory limitations of peripheral vision. Its distributed representation and image-computable nature also enable direct comparisons between model activations and neural activity in the brain. We show that after training to categorically search for objects in natural scenes, the model learns a general mechanism for search that generalizes to novel scenes and object categories, and exhibits human-like behavior while relying on neural representations analogous to those observed in the primate brain. Having a dynamic internal neural representation, the latent space of the model, provides us with the opportunity to not only study the brain-similarity of the learned representations but also to make testable predictions about the geometry and dynamics of these latent representations, the computations they support, and possible mechanisms that implement visual search behavior in the brain’s neural circuitry.

## Results

### Goal-driven modeling of visual search in natural scenes

To model goal-directed visual search in complex natural environments, we designed a behavioral task that follows the structure of human and animal psychophysics experiments. In each trial, the agent -here, our artificial neural network model-first views a frame containing an exemplar from a cued object category, the cue frame. This is followed by a natural scene image that contains an exemplar of the same category, the search frame. The agent begins fixating at the center of the scene and is then free to make a sequence of saccades to locate the target object (Fig. 1A). To train an artificial neural network to perform this task in a way that generalizes to novel images and objects, we constructed a large-scale natural scene search dataset. Specifically, we processed 10 million natural scene images from the Places365 dataset Zhou et al. (2017) using a state-of-the-art segmentation model (Mask R-CNN He et al. (2017)) to extract object masks for 80 categories (Fig. 1B). Each image yielded multiple search trials, depending on the number of object categories detected. For a given (image, cue category) pair, we generated a cue frame by randomly sampling an object exemplar from the target category and placing it at the center of a blank frame. This was followed by the corresponding natural scene image, which served as the search frame. Ground-truth target location masks were defined by overlaying a 10×10 spatial grid on the object segmentation mask, with each grid cell receiving a uniform probability if it was fully contained within the target object (Fig. 1B); Methods). We refer to this dataset as Places365-Search. In addition to training our model on this dataset, we evaluated its generalization using COCO-Search18 Chen et al. (2021), a benchmark dataset of human eye movements recorded during categorical search for objects in natural scenes, enabling systematic comparisons between the model’s search behavior and that of humans in novel scenes.

**Figure 1.**
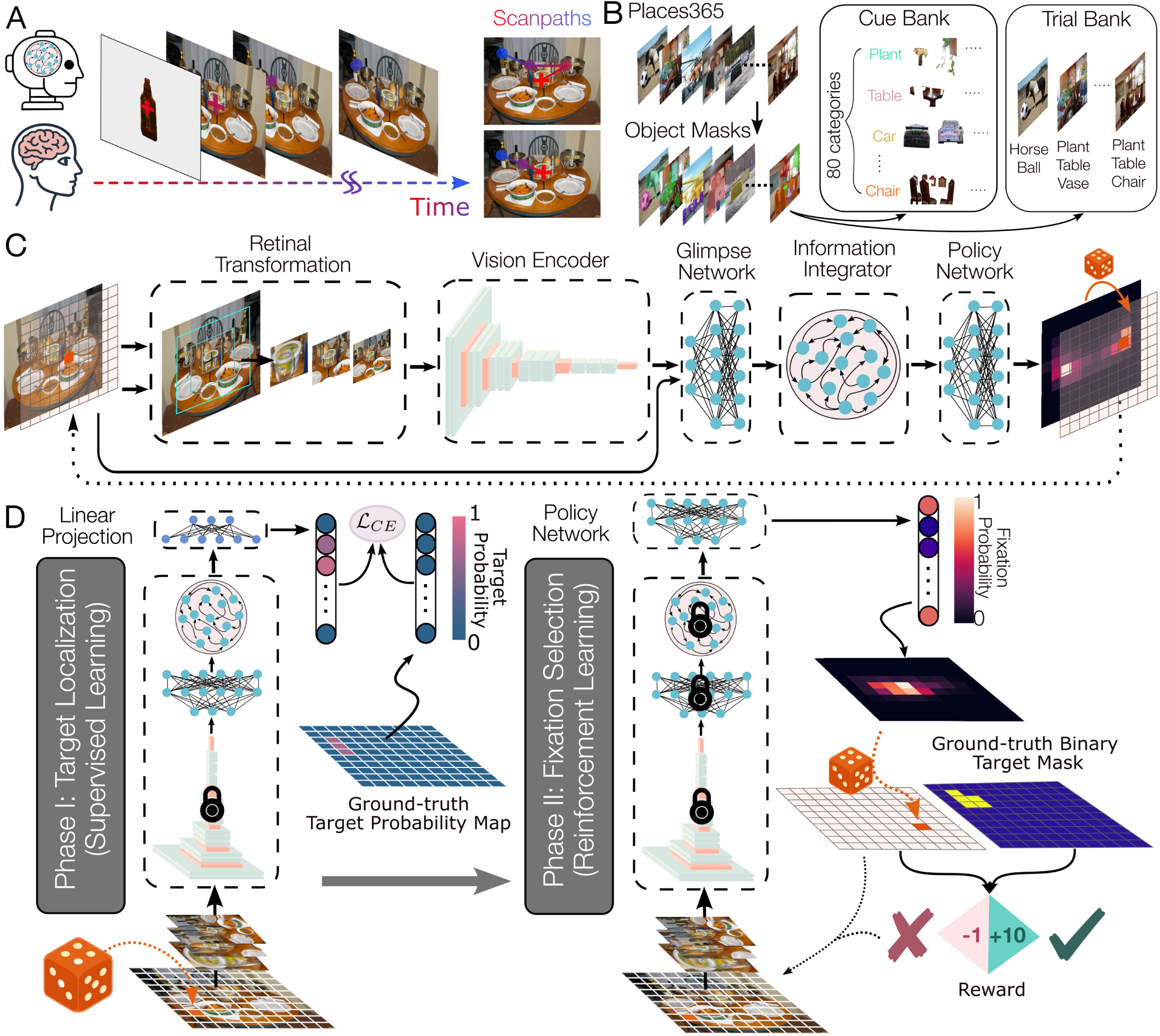
Behavioral paradigm and modeling framework. A. Behavioral paradigm. In each trial, the agent (human, monkey, or model) views a central cue depicting an object from the target category (e.g., a bottle). The subsequent search image is a natural scene containing another instance of the cued category. The agent freely shifts its gaze across the scene until the target is located or the trial ends. The resulting sequence of fixations, or *scanpath*, captures the spatiotemporal dynamics of overt visual search. **B. Task dataset generation pipeline.** 8,000,000 images of natural scenes from the Places365 dataset are passed through a state-of-the-art segmentation model to extract object masks for 80 categories (e.g., motorcycle, human, car, traffic light, kite). Extracted object instances were stored in a cue bank, serving as visual query examples for the corresponding cue categories. Each image and its annotated object categories formed the basis of the visual search trials. **C. Model architecture.** The model comprises several modules, each mimicking the role of different brain regions involved in overt visual search. A multi-resolution retinal sampling module approximates the progressive loss of visual acuity with increasing eccentricity. At each fixation, the retinal image is processed by a CNN pretrained on retinal ImageNet, producing a visual representation that is combined with fixation coordinates by the Glimpse Network. This representation is integrated over time by an Information Integration module (e.g., an LSTM), whose hidden state informs the Fixation Selector, a multilayer perceptron that outputs a probability distribution over possible next fixation locations. Together, these modules emulate the visual and fronto-parietal circuits guiding attention and gaze during search. **D. Model training.** Training proceeded in two phases. In Phase I (target localization), the Information Integration module and a linear readout were trained with supervised learning to predict the target probability map from a sequence of partial observations sampled at random fixation locations. In Phase II (fixation selection), the policy network was trained with reinforcement learning to generate a 10 × 10 fixation probability map from which the next fixation was sampled. The agent received a reward of +10 for fixating the target (i.e., hit) and a penalty of -1 for each non-hit step. During testing, given a cue and a search image, the trained model searched for the target instance through sequential fixations.

Our proposed model, the *Goal-directed Visual Search Model (GVSM)*, comprises four components (Fig. 1C):

1. **Visual encoder:** A convolutional neural network (CNN) pretrained on ImageNet for visual processing, which provides a biologically plausible approximation of the ventral visual stream Deng et al. (2009); Yamins et al. (2014); Ratan Murty et al. (2021); Bashivan et al. (2019); Cadieu et al. (2014); Khaligh-Razavi and Kriegeskorte (2014).
2. **Glimpse network:** Combines the retinal input with the current fixation location (efference copy), simulating peripheral encoding and gaze-contingent vision Mnih et al. (2014).
3. **Information Integrator:** An LSTM/transformer that integrates information across fixations, acting as an internal memory and taking on the role of the fronto-parietal attentional control network in guiding fixations, along with the fourth component Graves (2012); Vaswani et al. (2017).
4. **Policy network:** A multilayer perceptron (MLP) that generates a probability map over potential next fixation locations from the LSTM/transformer’s hidden state Rumelhart et al. (1986).

At each time step, the model receives a foveated visual input via a multi-resolution cropping mechanism that approximates eccentricity-dependent acuity Mnih et al. (2014). The CNN processes this input, and the resulting features are fused with the fixation vector through the glimpse network Mnih et al. (2014). This glimpse representation is then fed into the LSTM/transformer, which integrates across past fixations. The LSTM hidden state/most recent token from the transformer is passed to the policy network, which produces a 10×10 fixation probability map. The next fixation is sampled from this distribution. The trial continues until the model fixates the target (i.e., selects a grid cell within the ground-truth mask) or reaches a maximum step limit.

Model parameters were optimized using a two-stage training paradigm, inspired by prior work on saccade-augmented visual categorization Elsayed et al. (2019). In the first stage, we trained the Information Integrator (LSTM/transformer) via supervised learning to predict the target region from a random sequence of fixations over the image. In the second stage, we froze the LSTM/transformer weights and trained a policy network (MLP) using reinforcement learning (RL) to select fixation locations. During the supervised training stage, the LSTM/transformer learned to accumulate information across glimpses and build an internal representation of the target’s location. In the RL stage, the policy network leveraged this latent representation to guide efficient fixation selection. The agent was rewarded for locating the target and penalized for each unsuccessful fixation, thereby encouraging shorter search trajectories.

### GVSM replicates human search behavior

We evaluated GVSM’s generalization and behavioral alignment with humans by comparing it to three leading models of visual search. Two of these—IVSN and eccNET—are biologically inspired algorithmic models that compute a cue-similarity map over the visual scene using CNN-derived features. Fixations are selected using a winner-take-all strategy with inhibition of return. While IVSN assumes uniform resolution across the visual field, eccNET incorporates eccentricity-dependent pooling to mimic retinal transformations. As a third baseline, we included an inverse reinforcement learning (IRL) model trained on human eye-tracking data from COCO-Search18, serving as a gold-standard data-driven benchmark.

#### Generalization to novel scenes and objects

Humans can flexibly generalize their visual search behavior to any natural scene, containing diverse objects. A general model of visual search is naturally expected to not merely memorize training examples but to generalize to novel scenes and objects, reflecting the flexibility and robustness of human search behavior. We tested GVSM on a held-out dataset, the COCO-Search18 test set, which included novel cue exemplars and search scenes. GVSM achieved the highest overall accuracy (mean=85.67 ± S.D.= 0.87), significantly outperforming both the IRL model (*t* = 11.81, *p <* 10*^−^*^4^) and the algorithmic baselines (IVSN: *t* = 21.35, *p <* 10*^−^*^4^, eccNet: *t* = 88.19, *p <* 10*^−^*^4^; Fig. 2B), despite not being trained on eye-tracking data. This confirms that GVSM supports flexible, category-conditioned search in natural environments.

**Figure 2.**
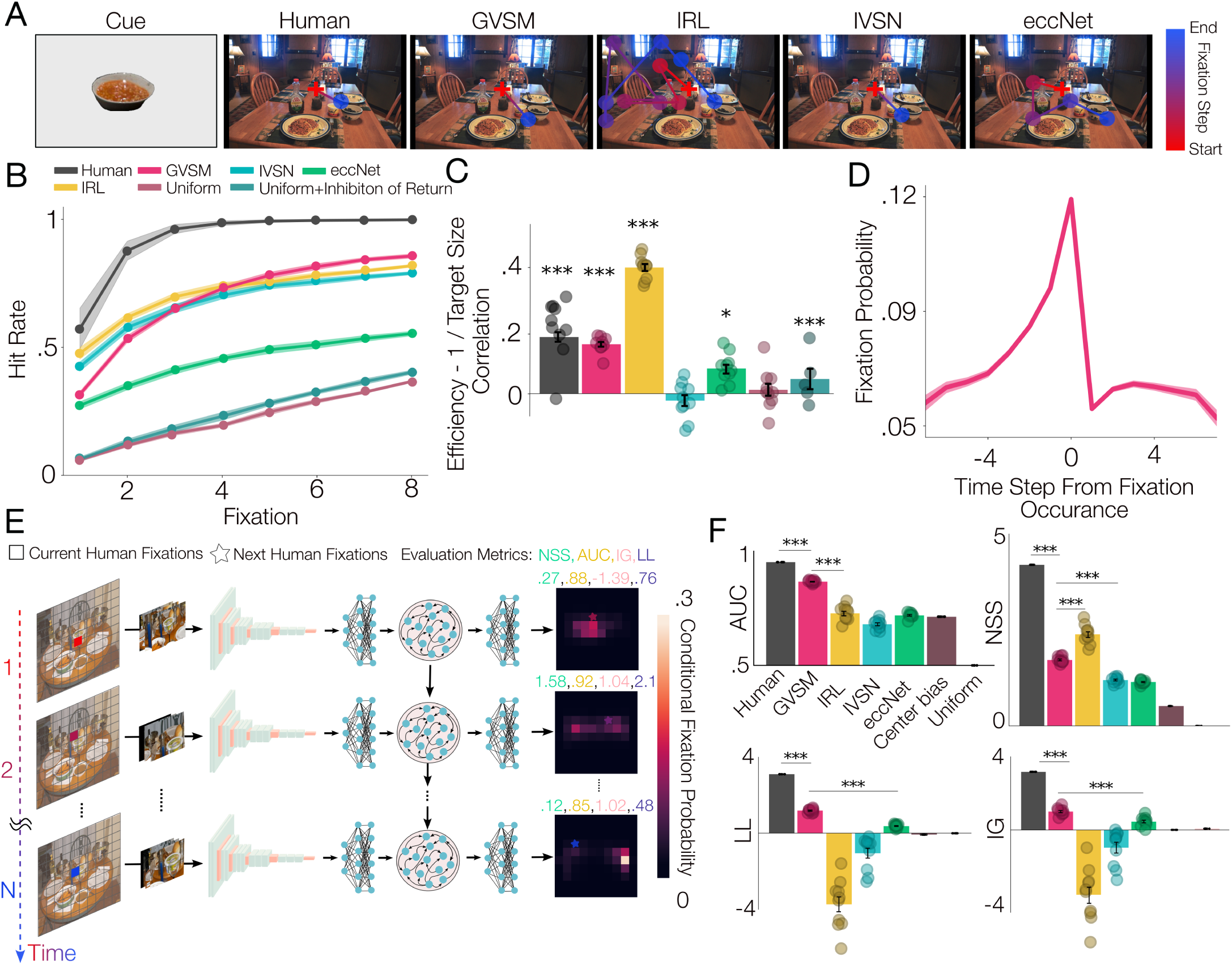
GVSM replicates human search behavior. A. Example test search scanpaths. Search scanpaths for a human subject, GVSM, and baseline models on an example trial from the COCO-Search18 test set. **B. Cumulative performance curves.** Hit rate, the proportion of trials in which the target was found, as a function of the number of fixations is shown. **C. Comparing the effect of target size on search efficiency across models.** Spearman correlation of the number of fixations required to find the target with the <u>^1^</u> averaged across participants or model instances. Each point represents a human participant, a run of the GVSM model, or a bootstrapped instance of the deterministic baselines. We bootstrapped the trials to create multiple instances of the deterministic baselines. Error bars represent the standard error magnitude (SEM) across the instances. GVSM exhibits the closest efficiency-target size relationship to humans (GVSM: Spearman’s *ρ* = 0.19, *p <* 10*^−^*^4^), Human: Spearman’s *ρ* = 0.17, *p <* 10*^−^*^4^). The stars indicate the significance of the correlation. Stars indicate significance from paired t-tests: *:*p <* 0.01, **:*p <* 0.001, ***:*p <* 10*^−^*^4^. **D. The evolution of fixation probability over time.** The probability of selecting each fixated location is plotted across time relative to when it is fixated. The x-axis denotes fixation offset (in time steps), and the y-axis indicates its probability in the fixation map. The solid line represents the average evolution curve across all the fixated locations in all the test trials, and the shaded areas represent their SEM. The probability of fixating at the location increases gradually, peaking at the fixation time step, then dropping to a nonzero value–showing that GVSM effectively acquires a human-like inhibition of return strategy without being explicitly instructed to do so. **E. Illustration of the fixation-by-fixation scanpath prediction method.** The model receives the sequence of human fixations, and at each time step, outputs a conditional fixation probability map (the fixation selector module’s output), *P* conditioned on the human fixation selection history thus far on the image *I*. Each conditional probability map is compared to the participant’s ground-truth fixation at that step using four standard metrics from the saliency prediction literature (Methods). **F. Trial-by-trial scanpath alignment.** GVSM outperforms all the baselines in terms of trial-by-trial alignment with human search scanpaths on three of the metrics (AUC: GVSM vs. IRL: *t* = 13.14, *p <* 10*^−^*^4^, LL: GVSM vs. eccNet: *t* = 19.85, *p <* 10*^−^*^4^, IG: GVSM vs. eccNET: *t* = 5.52, *p <* 10*^−^*^4^) and comes second on NSS, only to IRL (GVSM vs. IRL: *t* = −8.35, *p <* 10*^−^*^4^, GVSM vs. IVSN: *t* = 15.32, *p <* 10*^−^*^4^), which is a model trained on human eye-tracking data to mimic human behavior, thus serving as a reference here. Each dot represents a comparison of the model scanpaths with one subject, and error bars represent the SEM across those comparison pairs. Stars indicate significance level from paired t-tests as in A.

#### Human-like search efficiency

We next assessed how closely GVSM mirrored human search behavior. First, we analyzed search efficiency using cumulative performance curves (Fig. 2B), which quantify the proportion of successful trials as a function of fixation count. While all models underperformed humans, GVSM best approximated the human curve. To probe second-order structure in efficiency, we tested for sensitivity to target size. Humans required more fixations for smaller targets (Spearman’s mean *ρ* = 0.17 ± *SEM* = 0.01, *p <* 10*^−^*^4^; Fig. 2C), a pattern replicated by GVSM (Spearman’s mean *ρ* = 0.15 ± *SEM* = 0.01, *p <* 10*^−^*^4^; Fig. 2C). By contrast, IRL overestimated this effect (Spearman’s mean *ρ* = 0.37 ± *SEM* = 0.01, *p <* 10*^−^*^4^; Fig. 2C), and IVSN failed to capture it (Spearman’s mean *ρ* = −0.01 ± *SEM* = 0.01, *p* = 0.48; Fig. 2C). GVSM thus mirrors both aggregate efficiency and second-order dependencies of search efficiency on scene features (Fig. 2C).

#### Emergent inhibition of return

A hallmark of human search is inhibition of return (IOR): the tendency to avoid revisiting prior fixations with a high frequency Posner et al. (1985); Klein (1988, 2000). We asked whether GVSM exhibits similar dynamics. Using the Policy Network’s output probabilities, we tracked the likelihood of re-fixating previously visited locations over time. The probability peaked at fixation and declined thereafter (Fig. 2D), replicating the characteristic post-fixation suppression seen in human search. These findings reaffirm that fixation history plays a significant role in guiding eye movements Posner et al. (1985); Klein (1988, 2000); Smith and Henderson (2009), and demonstrate that GVSM captures this effect. By contrast, the baseline models rely on hand-engineered IOR mechanisms to prevent the model from repeatedly fixating the same location Zhang et al. (2018); Gupta et al. (2021); Zelinsky (2008); Pomplun and Gray (2007). While effective, this technical fix highlights a key limitation of such frameworks: they lack a dynamic internal representation that integrates information across fixations. Although engineering IOR is common even in data-driven models, growing evidence suggests that primates do revisit previously attended locations, albeit with lower probability Smith and Henderson (2009); Zhang et al. (2022); Wilming et al. (2013). GVSM’s ability to exhibit human-like inhibition of return without explicit behavioral constraints suggests that such dynamics can emerge naturally from optimizing task performance in a model that integrates partial observations over time.

#### Trial-by-trial scanpath consistency with humans

Finally, we tested fine-grained behavioral alignment between GVSM and humans by comparing full scanpaths on a fixation-by-fixation basis. At each step, we obtained the model’s the 10×10 fixation probability map conditioned on the subject’s prior saccades and evaluated it against the subject’s next fixation using standard saliency metrics Kümmerer and Bethge (2021) (Fig. 2E–F). GVSM achieved the highest scanpath similarity to human data, significantly outperforming eccNET and IVSN as well as IRL which is trained on human eye-tracking data to mimic human behavior, thus serving as a reference here (Fig. 2F). Taken together, these results show that GVSM captures both the coarse and fine structure of human visual search behavior, including generalization, efficiency, memory dynamics, and temporal fixation sequences, without access to human eye-tracking data.

### GVSM reproduces horizontal–vertical biases in human visual search

Human visual behavior is anisotropic across polar directions at isoeccentric locations, with visual performance typically enhanced along the horizontal axis compared to the vertical Barbot et al. (2021); Kupers et al. (2022); Chaikin et al. (1962); Carrasco et al. (2004); Corbett and Carrasco (2011); Mackeben (1999); Altpeter et al. (2000). This horizontal–vertical asymmetry extends to eye movements, where humans tend to fixate more frequently in horizontal directions Tatler and Vincent (2009); Najemnik and Geisler (2005); Koevoet et al. (2025); Bahill et al. (1975a). Consistent with prior work, we found that participants in the COCO-Search18 task exhibited a strong horizontal bias, with fixations most frequently aligned along the *x*-axis (Fig. 3A). GVSM captured this bias more faithfully than the baselines, demonstrated by a lower distributional mismatch from human behavior (GVSM vs. Human: *t* = 3.96, *p <* 0.01, GVSM vs. IRL: *t* = −6.34, *p <* 10*^−^*^4^, GVSM vs. IVSN: *t* = −4.74, *p <* 0.001; Fig. 3B).

**Figure 3.**
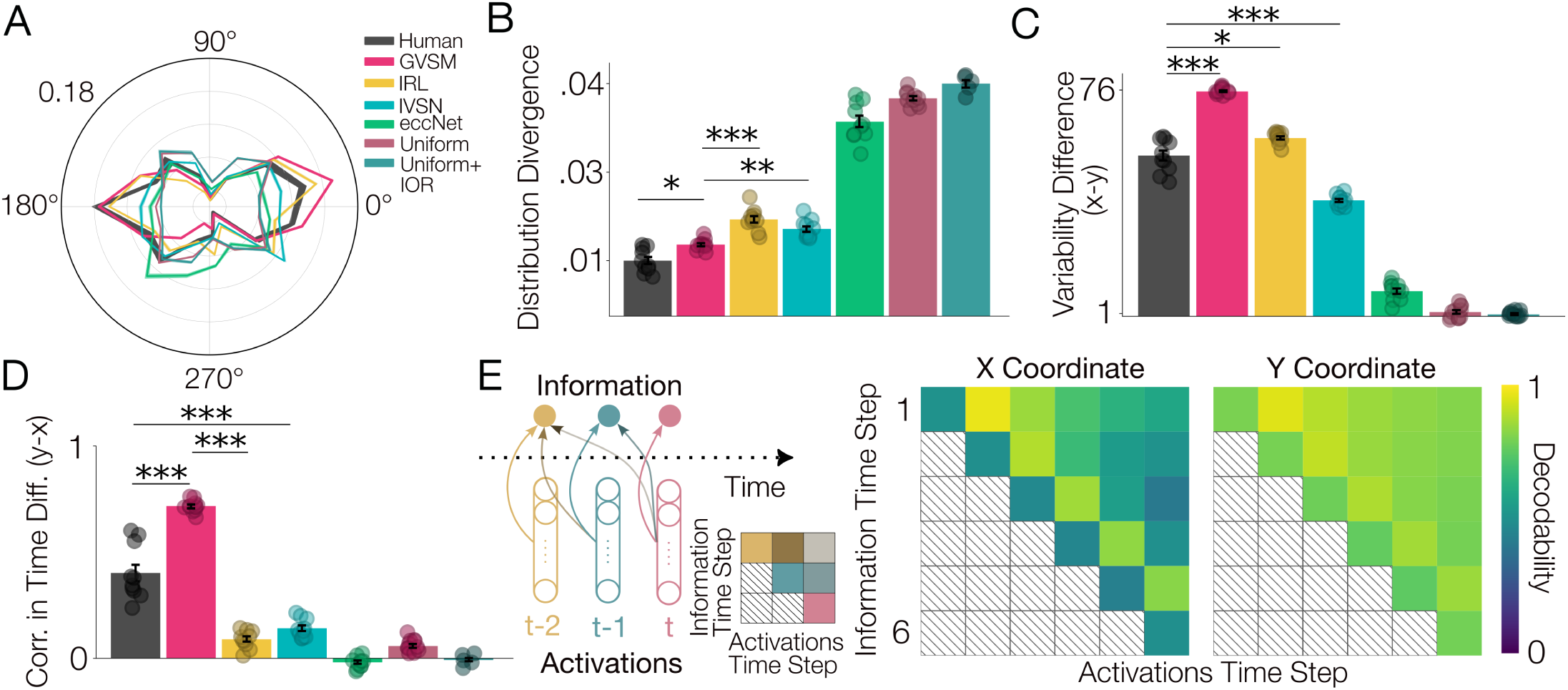
Human-like horizontal-vertical biases captured by GVSM. A. Histogram of fixation directions. Histograms of fixation directions show that both humans and GVSM exhibit a preference for horizontal over vertical fixations on the COCO-Search18 test set. Each curve represents the average probability distribution function of the fixation directions, averaged over model instances or human participants. Shaded bands represent the SEM across observer instances (human participants, GVSM runs, or the bootstrapped version of the deterministic baselines). Colors follow the same conventions as in panels B-D. Fixation directions are shown in degrees (0-360), and the range was divided into 15 bins. Each concentrated circle represents a frequency level (outermost circle=0.18). **B. Mismatch with the human distribution of fixation directions.** The distribution of GVSM’s fixation directions most closely matches that of human observers, i.e. its distribution of fixation directions are least different from those of humans (GVSM vs. Human: *t* = 3.96, *p <* 0.01, GVSM vs. IRL: *t* = −6.34, *p <* 10*^−^*^4^, GVSM vs. IVSN: *t* = −4.74, *p <* 0.001). The *y*-axis shows the distance between each model’s and humans’ distribution of fixation directions (the angle between the fixation location and the positive *x*-axis in the visual scene; Methods) averaged over instances (human participants, GVSM runs, or bootstrapped versions of the deterministic baselines). Each dot represents an instance, and the error bars represent the SEM across them. **C. Variability difference along the** *X* **and** *Y* **axes.** Standard deviation of fixated *X* coordinates minus that of the fixated *Y* coordinates is shown for all the models and humans, averaged across instances. Each point represents an instance and error bars represent the SEM across them. GVSM’s fixations are more variable across the *x*-axis compared to the *y*-axis, capturing the same horizontal biases as humans. Although the bias is stronger in GVSM compared to humans (GVSM vs. Humans: *t* = 12.12, *p <* 10*^−^*^4^, IRL vs. Humans: *t* = 3.52, *p <* 0.01, IVSN vs. Humans: *t* = −8.13, *p <* 10*^−^*^4^). **D. Difference in correlation across time along the** *X* **and** *Y* **axes.** Average correlation of the consecutive fixated *X* and *Y* coordinates across time, averaged over the pairs, is computed for all the instances of models and human participants. Each dot represents the difference between the average correlation across time along *X* axis minus that along the *Y* axis for an instance (a human participant, or a GVSM run, or a bootstrapped version of the baselines), and error bars represent the standard deviation across them. GVSM’s fixated *y* coordinates are more correlated across time compared to the *x* coordinates, although the bias is stronger than that of humans (GVSM vs. Humans: *t* = 7.56, *p <* 10*^−^*^4^, IRL vs. Humans: *t* = −6.67, *p <* 10*^−^*^4^, IVSN vs. Humans: *t* = −5.86, *p <* 10*^−^*^4^). **E. Memory encoding of previous fixations in the model’s latent space.** Left: Memory decoding analysis schematic. To quantify how information from past fixations is maintained over time, we trained linear regression models to decode specific variables (e.g., fixation coordinates) from the hidden state of the Information Integration module (LSTM). Each row of the resulting matrix indicates the time step when information became available, i.e. the Information Time Step, and each column shows when it is read out from the network’s activations, i.e., the Activations Time Step. Right: The decoding accuracy of linear regression models trained to decode fixated coordinates from the model’s latent space at the same time step (diagonal) or later time steps (off-diagonal) is shown. Each row represents the time step corresponding to the information to be decoded, here, the upcoming fixation location, and the columns represent the time step of the model’s latent space from which the information is decoded. Upcoming fixation locations are linearly decodable from the model’s latent space (*x* : 0.48 ± 0.03; *y* : 0.78 ± 0.02, shuffled chance level = 0). Previously fixated (*x, y*) locations are decodable from the unit activations of the model’s latent space at later time steps (off-diagonal elements) (*x* : 0.69 ± 0.16; *y* : 0.85 ± 0.04, shuffled chance level = 0). Notably, fixation selection history is more strongly decodable compared to the upcoming fixation coordinates (off-diagonal values are higher than the diagonal). Consistently, unlike the selection stage, *X* coordinates are reliably decoded, indicating specificity of the horizontal bias to selection rather than memory encoding.

A second signature of this asymmetry is the greater variability in fixated *x* -coordinates compared to *y* -coordinates. All models captured this effect, but the difference between *x* and *y* variability in GVSM was closest to human behavior (Fig. 3C). We further examined the temporal dependencies in fixation patterns, computing correlations across time for each spatial axis. In humans, fixated *y* -coordinates were strongly correlated over time, whereas *x* -coordinates were not (Fig. 3D), reflecting the greater temporal stability of vertical fixation locations. GVSM best reproduced this dissociation, resembling the observed human difference in correlation between axes.

We next asked whether these behavioral asymmetries were reflected in the model’s internal state. During inference, the latent space of GVSM—defined as the hidden state of the Information Integrator (LSTM or transformer)—can be viewed as a computational proxy for dynamic representations in the primate brain’s fronto-parietal attention network. To probe the encoding of upcoming fixation locations, we trained linear decoders to predict the next fixation’s *x* and *y* coordinates from the model’s latent state at each time step. Decoding accuracy was substantially lower for *x* than *y* coordinates (*x* : 0.48 ± 0.03; *y* : 0.78 ± 0.02, shuffled chance level = 0), revealing a neural correlate of the behavioral *x–y* asymmetry (Fig. 3E).

We then examined whether this asymmetry also extended to memory representations. Linear decoders trained to recover the previous fixation’s *x* and *y* coordinates from the latent space revealed high decodability for both axes (*x* : 0.69 ± 0.16; *y* : 0.85 ± 0.04, shuffled chance level = 0). However, these results had distinct correlational structures: while *y* coordinate memory could be partly explained by strong temporal autocorrelation (mean Spearman’s *ρ* = 0.60±*SEM* = 0.01, *t* = 78.4, *p <* 10*^−^*^4^), *x* coordinates showed negligible autocorrelation (mean Spearman’s *ρ* = –0.1±*SEM* = 0.01, *t* = –13.08, *p <* 10*^−^*^4^; Fig. 3D), suggesting that the memory trace for horizontal locations was not simply due to temporal persistence. These findings indicate that GVSM stores horizontal and vertical fixation history with comparable fidelity, despite greater behavioral variability along the horizontal axis.

Notably, GVSM captured these human-like asymmetries despite lacking any built-in oculomotor constraints—suggesting that such biases may emerge from task structure and visual scene statistics alone, rather than from low-level motor priors Tatler and Vincent (2009); Torralba and Oliva (2003). Supporting this interpretation, GVSM also reproduced other canonical properties of human saccades, including the skew toward shorter amplitudes and the tendency for horizontal saccades to be longer than vertical ones Tatler and Vincent (2009); Bahill et al. (1975b); Gajewski et al. (2004); Pelz and Canosa (2001); Tatler et al. (2006).

### GVSM reproduces canonical neural motifs of primate visual search

To test whether GVSM captures the neural phenomena traditionally implicated in visual search, we analyzed its latent representations using paradigms closely aligned with primate neurophysiology experiments. We first introduced a delay period between the cue and search frames, mirroring behavioral protocols used to study working memory in fronto-parietal areas such as FEF, LIP and VPA Bichot et al. (2015). We also ran simplified detection trials in which a single target appeared in one of nine possible grid locations, allowing us to map each unit’s receptive field (RF) and quantify stimulus selectivity (Methods).

#### Persistent activity and cue-category coding

Category-selective units in GVSM exhibited strong persistent activity during the delay period: they responded more strongly to their preferred cue category compared to non-preferred categories, analogous to the sustained encoding observed in VPA neurons Bichot et al. (2015) (Fig. 4A). At the population level, we identified the *encoding subspace* for cue category by fitting linear support vector machines (SVMs) to decode category identity from latent activity at each time step (chance level=1/18=0.05). Cue category was reliably decodable throughout the trial (diagonal; Fig. 4B). Moreover, cross-temporal generalization analyses revealed that decoders trained at one time step generalized across other time steps (off-diagonal; Fig.4B), indicating a stable cue-category code in GVSM’s latent space. This result predicts that the primate fronto-parietal network maintains a similarly stable cue-category representation across time during visual search—a hypothesis that can be experimentally tested.

**Figure 4.**
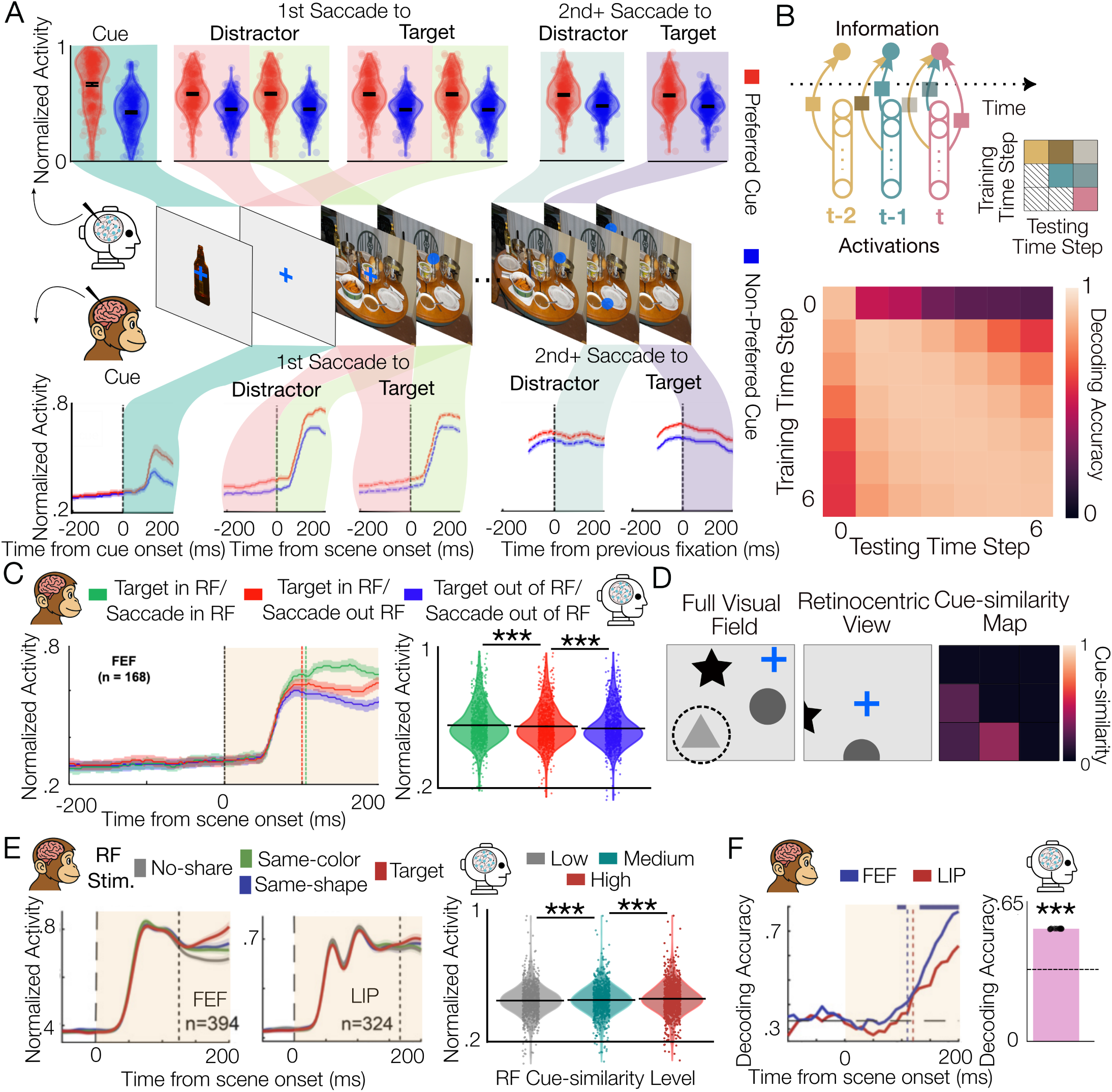
Brain-like cue-similarity representations in GVSM latent space. A. Persistent activity and cue-category coding. Responses of category-selective units when their preferred category was the cue (red) versus a non-preferred category (blue). Category-selective units responded more strongly when the cue category matched their preferred stimulus, exhibiting persistent cue-selective activity across the trial. Unit responses are shown (top: model latent units; bottom: reproduced from Bichot et al. (2015), VPA neurons in macaque monkeys) throughout the trial: from left to right: after cue-frame presentation, preceding the first saccade to the target or a distractor: after the delay frame (red shading) and after the scene onset (light green shading), and preceding subsequent saccades to the target (purple shading) or a distractor (blue shading). Shaded areas specify the period of time to which the activity plots correspond. Black dots indicate medians. **B. Cue-category encoding subspace.** Top: schematic of the cross-time generalization decoding analysis. Bottom: heatmap of decoding accuracy across training and testing time steps. Linear classifiers (SVCs) trained to decode the cue category from the model’s latent space reliably decoded the cue category at the training time step (diagonal). Off-diagonal generalization accuracy indicates that the same encoding subspace supports cue-category representations across time, revealing a temporally stable code. **C. Spatial and feature selection effects.** Left: population response time courses from primate FEF neurons across three conditions (green: target in RF and saccade into RF; red: target in RF and saccade out of RF; blue: target out of RF and saccade out of RF), reproduced from Bichot et al. (2015). The shaded region (light orange) indicates the time window corresponding to the model’s time step. Right: responses of GVSM units in the same three conditions (black horizontal lines: median, dots: individual units). Units exhibited significant spatial selection effects (green vs. red; *t* = 8.62, *p <* 10*^−^*^4^, Cohen’s *d* = 0.06) and feature selection effects (red vs. blue; *t* = 9.68, *p <* 10*^−^*^4^, Cohen’s *d* = 0.11), paralleling the response patterns observed in primate fronto-parietal neurons. The shaded area specifies the period of time to which the model’s plot corresponds. **D. Retinocentric cue-similarity map computation.** Schematic illustrating how cue-similarity maps are computed from the model’s visual representations. Left: the full visual field, with the current fixation location indicated by the blue cross and the target (dashed circle) among distractors. Middle: the retinocentric view aligned to the current fixation. Right: the resulting 3×3 retinocentric cue-similarity map, computed by convolving the CNN features of the cue with those of the search frame. Warmer colors indicate higher similarity between the local search image region and the cue exemplar. **E. Neural responses scale with cue-similarity.** Left: normalized firing rates of neurons in primate FEF (left panel) and LIP (middle panel) from Sapountzis et al. (2018), showing stronger responses to stimuli with higher cue-similarity. Curves are color-coded based on cue-similarity of the RF stimulus (red: target, green: distractor sharing the target’s color, blue: distractor sharing the target’s shape, and grey: distractor not sharing features with the target). Right: GVSM unit activity shows the same graded response pattern, with significantly higher normalized activity for stimuli with increasing cue-similarity levels (grey: low, teal: medium, and red: high; low vs. medium: *t* = −6.33, *p <* 10*^−^*^4^, Cohen’s *d* = −0.05, medium vs. high: *t* = −9.46, *p <* 10*^−^*^4^, Cohen’s *d* = −0.08) (black horizontal lines: medians; dots: individual units). The shaded area specifies the period of time to which the model’s plot corresponds. **F. Cue-similarity level decoding.** Left: time-resolved decoding accuracy of cue-similarity levels from population activity in primate FEF (blue) and LIP (red), reproduced from Sapountzis et al. (2018). Decoding accuracy rises above chance shortly after array onset, indicating that these areas encode cue-similarity during search. Right: GVSM latent-space decoding accuracy for the same 3-way classification task (low-mid-high cue similarity). We trained nine separate linear decoders (one per location in the 3×3 grid; D) and averaged their decoding accuracies. Average decoding accuracy is significantly above chance (Mean decoding accuracy = 0.55, Shuffled decoding accuracy = 0.33, *p <* 10*^−^*^4^), demonstrating that the model’s population activity reliably encodes cue-similarity levels. (black horizontal line: chancel level=0.33; dots: splits). The shaded area specifies the period of time to which the model’s plot corresponds.

#### Spatial and feature selection effects

Another hallmark of the fronto-parietal areas (VPA, FEF, LIP) is the modulation of neural responses by both spatial and feature selection Bichot et al. (2015); Sapountzis et al. (2018). Following the methodology in Bichot et al. (2015), we measured unit responses in three conditions: (1) the target was inside the RF and the first saccade was directed to it, (2) the target was inside the RF but the saccade was directed elsewhere, and (3) the target was outside the RF and the saccade was directed elsewhere. Units in GVSM showed significantly stronger responses when the saccade was directed into the RF (spatial selection) and when the target was present inside the RF independent of the saccade (feature selection) (stats; Fig. 4C), closely paralleling effects reported in primate physiology Bichot et al. (2015).

#### Cue-similarity encoding

The feature selection effect described above has been interpreted as evidence that fronto-parietal neurons encode retinocentric *cue-similarity maps* Bichot et al. (2015); Sapountzis et al. (2018); Bisley and Mirpour (2019); Egner et al. (2008); Thompson and Bichot (2005); Wardak et al. (2004); McPeek and Keller (2002); Nobre (2001); Kusunoki et al. (2000). In Sapountzis et al. (2018), cue-similarity was measured at a more granular level by having distractors with a gradient of cue-similarity from no shared features with the cue to sharing the shape or color with the cue. To probe this in GVSM, we computed cue-similarity over the search frame by convolving CNN feature maps from the cue and search frames, yielding a 3×3 similarity map (Methods, cue-similarity map computation; Fig. 4D). GVSM units produced the graded modulation observed in FEF and LIP, responding more strongly when their RF contained higher cue-similarity stimuli(Fig. 4E). Cue-similarity level could be decoded at the population level similar to FEF and LIP. Linear SVMs trained to classify high, medium, and low cue-similarity reliably decoded similarity level from GVSM’s latent activity (Mean decoding accuracy = 0.55, Shuffled decoding accuracy = 0.33, *p <* 10*^−^*^4^; Fig. 4F). Overall, these findings indicate that a cue-similarity map representation akin to those observed in the fronto-parietal areas such as FEF and LIP emerges in the latent space of the model trained to do visual search.

### Emergent neural correlates of saccade goal selection

Having shown that GVSM replicates neural signatures of cue-category memory and attentional modulation observed in fronto-parietal areas such as VPA, FEF, and LIP, we next asked whether similar correlates of saccade goal selection emerge in the model’s latent representations. Specifically, we tested whether GVSM units exhibit activity patterns resembling those found in primate fronto-parietal and midbrain regions, including FEF, LIP, and the superior colliculus (SC) that support the planning and selection of saccades.

#### Encoding of upcoming fixation goals

Following the methodology of Sapountzis et al. (2018), we first compared unit responses immediately prior to the model’s first fixation, contrasting trials where the saccade was directed into a unit’s RF versus away from it—regardless of the cue-similarity at that location. Similar to neurons in FEF and LIP, GVSM units responded more strongly when the upcoming fixation targeted their RF (Fig. 5A). This modulation was also evident at the population level: a binary linear classifier trained to decode whether the next fixation would land within the RF achieved high accuracy (0.75 ± 0.002; Chance: 0.5 ± 0.002; Fig. 5B), indicating that the model’s latent activity reliably encodes information about saccade goals.

**Figure 5.**
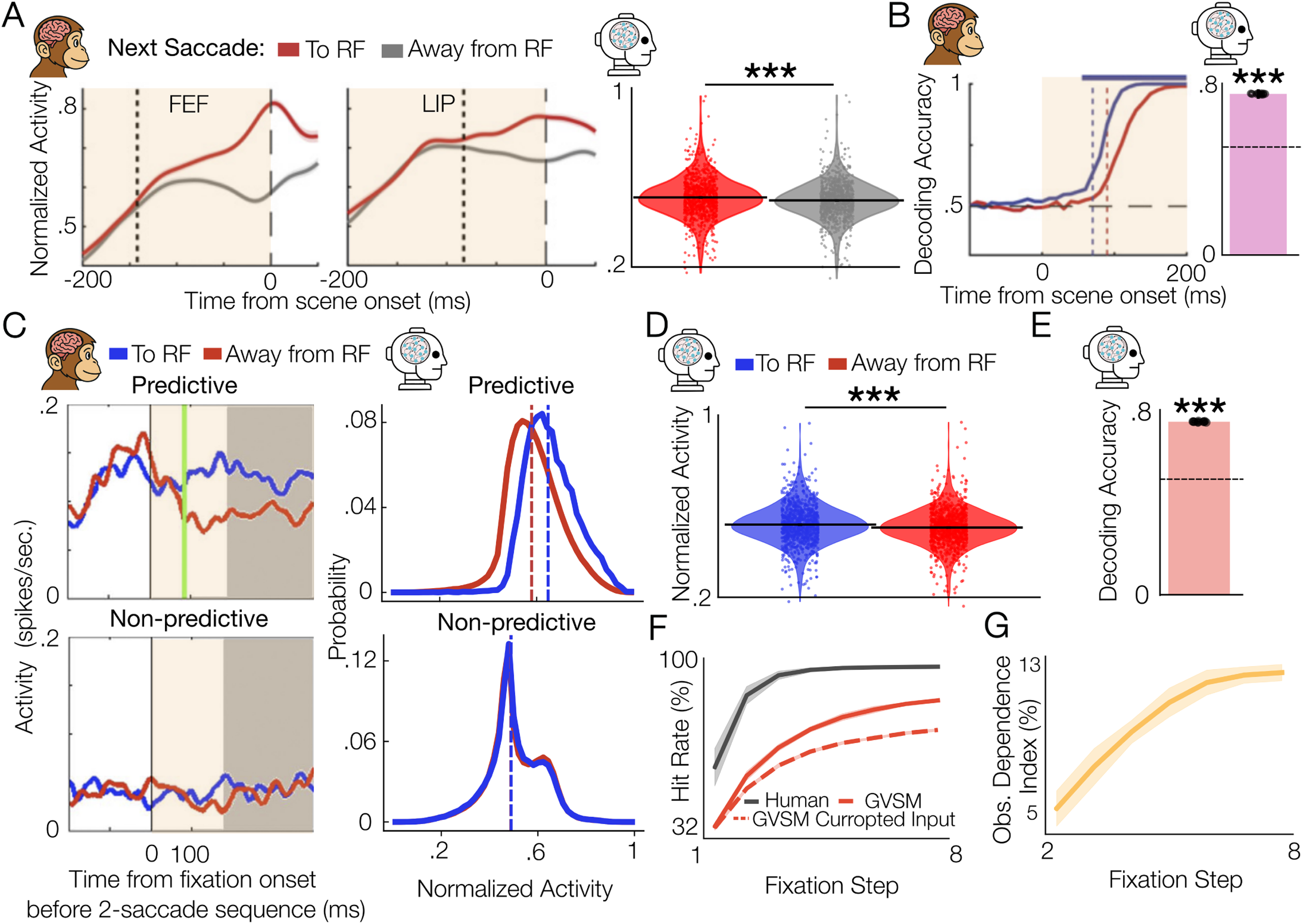
Emergent neural correlates of saccade goal selection. A. Population responses in primate and model reflect encoding of the upcoming saccade target. Left: Normalized firing rates of neurons in primate FEF and LIP (reproduced from Sapountzis et al. (2018)), aligned to the onset of the first saccade. Responses are shown for trials in which the saccade landed within the neuron’s RF (red) versus outside it (gray). Right: GVSM latent unit activity shows a similar effect. Units exhibited significantly higher pre-saccadic responses when the first fixation was directed into their RF (*t* = 12.62, *p <* 10*^−^*^4^, Cohen’s *d* = 0.16), indicating that the model encodes information about the upcoming saccade goal. Black horizontal lines indicate medians; colored dots indicate individual units. The shaded area specifies the period of time to which the model plot corresponds to. **B. Decoding of upcoming saccade target from population activity in primate and model.** Left: Time-resolved decoding accuracy of whether the first saccade was directed into a neuron’s RF, based on population activity in FEF (blue) and LIP (red), reproduced from Sapountzis et al. (2018). Decoding accuracy rises sharply following array onset, indicating that these areas encode the upcoming fixation target. Right: GVSM latent activity at the first time step supports similarly robust decoding of the upcoming saccade direction. A binary linear classifier trained to predict whether the first fixation landed within the unit’s RF achieved significantly above-chance accuracy (0.75 ± 0.002; Shuffled: 0.5 ± 0.002).The shaded area specifies the period of time to which the model’s plot corresponds. The dashed line indicates the chance level, and the black dots indicate individual splits. **C. Activity predictive of the goal of the second future saccade.** Top: A subset of macaque FEF and model units exhibit predictive activity for saccade targets two steps ahead. (left) Example FEF neuron from Phillips and Segraves (2010) showing elevated activity when the second upcoming fixation will land within its RF (blue: to RF; red: away), indicating predictive coding of future saccade goals. (right) Example GVSM “future-predictive” units show a similar pattern. Distributions of an example model unit responses prior to the first saccade are plotted separately based on whether the second upcoming fixation is to the RF (blue) or not (red). These units responded more strongly when a future fixation would target their RF, despite no stimulus yet being present there (Mann–Whitney U test: *U* = 6.24 ∗ 10^9^, *p <* 10*^−^*^4^, Cliff’s *δ* = 0.37). Bottom: A subset of FEF and model units do not exhibit predictive activity for saccade targets two steps ahead. (left) Example non-predictive neuron from Phillips and Segraves (2010) whose activity does not vary with the second upcoming fixation target. (right) Model units classified as non-predictive show no significant difference in pre-saccadic activity between trials where the second fixation is to the RF (blue) or not (red), indicating a lack of future-goal sensitivity (right; example unit: Mann–Whitney U test: *U* = 3.19 ∗ 10^10^, *p >* 0.95, Cliff’s *δ* = 0.00). **D. Latent activity is modulated by future fixation targets.** Normalized activity of GVSM latent units at the first time step, grouped by whether the second upcoming fixation would land within the unit’s RF. Units exhibited significantly higher responses when a future fixation was directed to their RF (*t* = 12.63, *p <* 10*^−^*^4^, Cohen’s *d* = 0.17), indicating that population activity carries information about saccade targets beyond the immediate next move. Horizontal black lines indicate means; colored dots represent individual units. **E. Decoding the second upcoming fixation from the latent state.** A binary linear classifier trained to predict whether the second upcoming fixation would land in the RF, based on latent activity at the first time step, achieved significantly above-chance accuracy (0.75 ± 0.003; Shuffled: 0.5 ± 0.003). This confirms that GVSM encodes multi-step saccade plans in its internal representations. The black dashed line indicates the chance level; black dots represent accuracy for different cross-validation splits. **F. GVSM maintains performance despite the removal of visual input after the first fixation.** Hit rate as a function of fixation step for humans (black), GVSM (solid red), and an ablated version of GVSM (dashed red) that received the search frame only once at the first time step and blacked-out input thereafter. Despite the lack of visual information beyond the first step, the ablated model achieved substantial accuracy, suggesting that its fixation policy was partially informed by the initial glimpse. **G. Later fixations depend more on updated visual input.** Behavioral input-dependence index across fixation steps, quantifying the performance drop caused by ablating visual input after the first glimpse. The increasing index with fixation number indicates that early fixations rely more heavily on preplanned trajectories, while later fixations increasingly depend on updated sensory input.

#### Predictive activity for future fixations

Neurons in FEF and the superior colliculus (SC) exhibit predictive signals for saccades multiple steps into the future Phillips and Segraves (2010); Shen and Paŕe (2014). To assess whether similar multi-step predictive activity emerges in GVSM, we analyzed model units based on whether their activity predicted fixation goals two steps ahead. Echoing primate results, we identified two functional classes of units: those showing elevated activity when the second upcoming fixation was to their RF: *predictive units* (Fig. 5C, top), and those that did not: *non-predictive units* (Fig. 5C, bottom). At the population level, GVSM activations preceding the first saccade were significantly modulated by whether the second upcoming fixation would land within the RF (Fig. 5D), and a binary classifier trained to decode this variable achieved high accuracy (0.75 ± 0.003; Chance: 0.5 ± 0.003; Fig. 5E). These findings suggest that GVSM encodes long-range saccade plans within its latent state.

#### Behavioral evidence for multi-step planning

If GVSM indeed plans multiple fixations in advance, then disrupting later sensory input should not immediately abolish performance. To test this, we conducted an ablation study where all visual input after the first step was replaced with black frames. Despite this manipulation, GVSM maintained substantial accuracy, indicating that its fixation policy had already committed to a partially preplanned trajectory based on the initial glimpse (Fig. 5F). The performance decline increased with longer trial lengths (i.e., more fixations), suggesting that later stages of search behavior rely increasingly on updated visual input (Fig. 5G).

Together, these results show that GVSM exhibits predictive dynamics strikingly similar to those observed in primate fronto-parietal and subcortical regions involved in saccade planning. Model units encode not only the immediate saccade target but also longer-horizon fixation plans, mirroring the forward-looking representations documented in FEF and SC. These emergent dynamics highlight GVSM’s potential as a computational model of visual search that captures both the behavioral structure and internal planning signals observed in the brain.

### Geometry and temporal structure of the latent cue-similarity representation

Having shown that unit activations in GVSM’s Information Integrator are modulated by cue-similarity in their RFs, mirroring cue-similarity selectivity observed in areas such as VPA, FEF, and LIP, we next examined the representational geometry of this encoding and how it evolves across time. Previously, we demonstrated that GVSM encodes a distributed retinocentric cue-similarity map at the population level at the first time step, analogous to cue-similarity map represen-tations found in fronto-parietal areas. Here, we extended that analysis by training separate linear regression decoders to predict cue-similarity values from latent activity at each time step (Fig. 6A). Decoding performance remained consistently high throughout the trial (Fig. 6B), indicating that the cue-similarity map is maintained in the model’s latent space across time.

**Figure 6.**
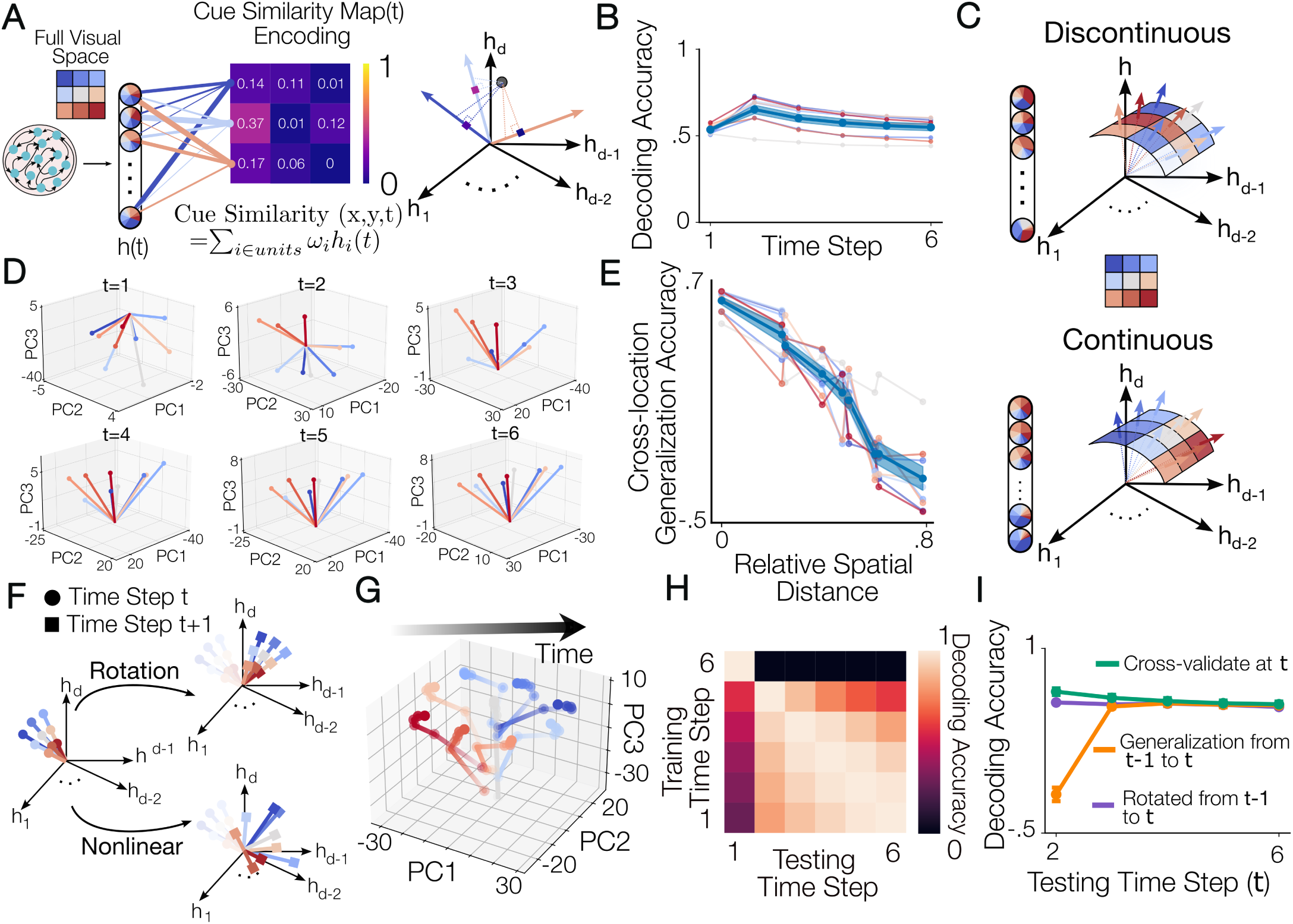
Geometry and temporal structure of the latent cue-similarity representation. A. Cue-similarity map encoding analysis schematic. Left: Activity patterns of the units in the Information Integration module (the recurrent network) were treated as latent neural population responses analogous to those observed in fronto-parietal attention control areas such as FEF and LIP. To test whether these units encode cue-similarity maps across the visual field, we trained linear regression models to decode the cue-similarity value at each spatial location from the population activity vector **h**(*t*). In the schematic, each unit contributes to the encoding of cue-similarity at different locations (illustrated by pinwheel diagrams color-coded by visual-field position). The color of each projection line outgoing from the units to the cue-similarity map schematic indicates the target location in the visual field (3 example locations shown), and its thickness reflects the relative weight of that unit’s contribution to the encoding of cue-similarity at that location. Right: An example schematic cue-similarity map encoded in the model’s latent space is shown. The latent state **h**(*t*) at each time step (black dot) can be represented in a high-dimensional state space, where each axis corresponds to the activation of one unit. Decoding axes (colored arrows) correspond to the learned weight vectors for three example spatial locations. The projection of **h**(*t*) onto each axis yields the cue-similarity value at the corresponding location. **B. Cue-similarity map encoding in the model’s latent space**. Decoding accuracy of cue-similarity values from GVSM latent activity at each time step using separately trained linear regression decoders as demonstrated in panel A, across time. Decoding remained consistently above chance across the trial, indicating that the cue-similarity map is preserved in the model’s internal representation over time. Error bars indicate SEM across nine spatial locations. **C. Schematic of potential geometries of the latent cue-similarity map representation.** The model could encode cue-similarity in either a discontinuous (top) or continuous (bottom) representational geometry. In a discontinuous representation, each visual location is encoded in an independent subspace, regardless of spatial proximity (top). The colored pinwheels indicate each unit’s relative contribution to encoding cue-similarity at different locations in the visual field. In this configuration, a unit may contribute strongly to distant locations with no spatial organization across units. In a continuous representation, cue-similarity at nearby locations are encoded by more aligned subspaces in the model’s latent space (bottom). Here, each unit’s contribution to cue-similarity varies gradually with spatial distance, resulting in more homogeneous color patterns across pinwheels. **D. Geometry of the latent cue-similarity map representation in the PC space.** Cue-similarity decoders are projected onto a shared 3D principal component space and color-coded by visual field location. Axes corresponding to spatially adjacent locations are more closely aligned, while those for distant locations diverge—revealing a continuous topographic structure in the latent representation. **E. Cross-location generalization supports a continuous topographic code.** Generalization accuracy of linear regression decoders trained on cue-similarity at one location and tested at others decreases as relative spatial distance between the locations increases, indicating that nearby locations share more overlapping encoding subspaces than distant ones, consistent with a continuous, topographically organized representation. **F. Schematic of possible transformations of the cue-similarity encoding subspace over time.** Illustration of hypothetical geometric relationships between cue-similarity representations at consecutive time steps. The encoding subspace may remain stable, rotate, or undergo a nonlinear transformation as the trial progresses. Each colored vector represents a decoder for a specific spatial location at time step *t* (circles) or *t* + 1 (squares). **G. PCA visualization of decoder trajectories reveals stabilization of the encoding subspace.** Decoder axes trained to predict cue-similarity at each time step were projected into a shared 3D principal component space. Trajectories are color-coded by spatial location (inset in panel A), and each dot represents a decoder from a specific time step, with darker points indicating later time steps. Decoder directions initially diverge from their configuration at time step 1 and gradually converge to a stable arrangement. Trajectories for the same spatial location remain tightly clustered, while those for different locations separate, reflecting stable and spatially distinct encoding across time. **H. Cross-time generalization of cue-similarity decoding.** Decoding accuracy of linear regressors trained at one time step (row) and tested at the others (column). Generalization was low between the first and later time steps, but high across later steps, indicating that the cue-similarity encoding subspace undergoes a transformation after the initial glimpse and then stabilizes. **I. Procrustes analysis reveals rotational transformation of the encoding subspace.** Decoding accuracy for cue-similarity at each time step using decoders trained and tested on the same time point (green), decoders trained at *t*−1 and tested at *t* without alignment (orange), and decoders trained at *t*−1 rotated using an orthogonal Procrustes transform, and tested at *t* (purple). Rotated decoders recovered high accuracy, indicating that the transition from the first to the second time step reflects a geometric rotation of the encoding subspace.

#### Topographic organization of the cue-similarity map

We next investigated how the population encodes cue-similarity values across the visual field. Specifically, we compared two hypotheses: (1) a discontinuous representation in which similarity at each location is encoded independently, and (2) a continuous topographic organization in which spatially adjacent locations are encoded by overlapping subspaces (Fig. 6C).

To distinguish between these possibilities, we projected the learned decoders into a shared three-dimensional PC space.The result revealed a smooth, topographic structure (Fig. 6D): decoders corresponding to neighboring spatial locations oc-cupied nearby directions, while those for distant locations diverged. This topographic gradient suggests a continuous organization of retinocentric space within GVSM’s latent representation.

We then asked whether cross-location generalization supports this continuity. Indeed, decoder performance declined systematically as the distance between training and test locations increased (Fig. 6E), consistent with partially overlapping population codes for nearby positions and opposing codes for distant ones. In practice, this opposing embedding of cue-similarity for distant locations may be a mechanism of inhibition similar to that observed in PFC neurons with opposing ”memory fields” Funahashi et al. (1989); Goldman-Rakic (1995) Together, these findings show that GVSM not only encodes a retinocentric cue-similarity map but does so in a geometrically structured and topographically meaningful manner.

#### Transformation of the cue-similarity encoding subspace over time

We next asked whether cue-similarity values are encoded in a consistent subspace across time or whether the representation undergoes geometric transformation. The evolution of the encoding subspace can take one of three forms: it may remain stable, undergo a simple rotation, or change via a nonlinear transformation (Fig. 6F). To visualize this transformation, we applied PCA to the decoder axes across all time points. In the reduced space, the cue-similarity decoders rotated from their initial configuration at time step 1 and converged to a stable configuration across later steps (Fig. 6G). Notably, decoder trajectories corresponding to the same spatial location remained close to one another, whereas those for different locations diverged, suggesting that spatial information is preserved and stabilized over time (Fig. 6G). To test this observation, we first quantified the consistency of the cue-similarity map encoding subspace across time by evaluating the cross-time generalization of its decoders: for each time step, we trained decoders on latent activity and tested them at all other time steps (Fig. 6H). Decoders trained on the first time step generalized poorly to later time points, while decoders trained after the first step generalized well to one another (Fig. 6H). This indicates that the cue-similarity encoding undergoes a qualitative shift after the initial glimpse, then stabilizes, supporting the observed convergence towards a stable representation in the PC space. Next, we used Procrustes analysis to test whether the transformation at early time steps is rotational. Specifically, we learned an orthogonal transformation that best aligned the decoder axes from time *t* to time *t* + 1, using a subset of spatial locations for fitting the rotation transformation and evaluating the decoders of the held-out locations after transformation by that learned rotation on time *t* + 1 (Methods: Procrustes analysis). The high decoding accuracy of rotated decoders (Fig. 6I) suggests that the transition from time step 1 to 2 primarily reflects a rotation of the encoding subspace. After this rotation, the encoding remained stable, consistent with the cross-time generalization results.

### Neural mechanisms of maintenance and retrieval of memory during visual search

So far, we have shown behavioral and neural signatures consistent with those of primates in GVSM, with a retinocentric cue-similarity map and memory playing important roles in shaping these behaviors. 1) We showed that the model units encode a retinocentric cue-similarity map similar to those encoded by VPA, FEF, and LIP in the primate brain presumed to guide fixation selection in visual search. 2) We showed that cue category is consistently encoded in the same stable subspace across time (Fig. 4 A-B), consistent with this code in the VPA neurons Bichot et al. (2015). 3) the model exhibits a human-like inhibition of return characteristics with a low probability of revisiting locations (Fig. 2D). This requires the model to have access to the information of prior fixations, i.e,. fixation selection history, which we found to be preserved in the model’s latent space by showing that previously fixated coordinates could be linearly decoded from the model’s latent space (Fig. 3F). 4) and activity predictive of fixations multi-steps into the future, similar to those of FEF neurons (Fig. 5C-E), with performance being largely preserved even with ablating the visual input after the first presentation of the search frame (Fig. 5F-G). This suggests the presence of a memory of the past cue-similarity maps in the model’s latent space, affecting fixation selection (Fig. 5G-H). Therefore, the current retinocentric cue-similarity map computed from the memory of cue-category representation and the incoming retinocentric glimpse of the search frame, the memory of past observations’ cue-similarity maps, and the fixation selection history, are of representations guiding selections in the model, consistent with the primates (Fig. 7A) Bichot et al. (2015); Bisley and Mirpour (2019); Sapountzis et al. (2018); Klein (2000); Torbaghan et al. (2012); Dorris et al. (2002); Zelinsky and Bisley (2015). Therefore, to accurately perform the visual search task, GVSM must retain crucial information about its prior observations and actions. Here, we asked how the model retains and reuses the memory of the cue-similarity map and fixation selection history across time. We reasoned that the Information Integrator component of GVSM may implement one of two distinct mechanisms to store memory representations of retinocentric cue-similarity maps or fixation selection history (Fig. 7B).

- **H1: Slot-based memory subspaces Whittington et al. (2025); Wang et al. (2025); Luck and Vogel (1997)**: In this framework, the model’s latent space is partitioned into distinct, time-indexed subspaces, or “slots”, with each slot dedicated to storing a single item, such as a computed cue-similarity map or a visited fixation location. Once an item is encoded, it remains in its designated slot until retrieval. These slots occupy unique and temporally stable regions of the latent space, supporting sustained, interference-resistant memory representations over time (Fig. 7B; left).
- **H2: Chronologically indexed memory subspaces Lei et al. (2024)**: In this alternative framework, memory is organized by the relative age of encoded content. The latent space is divided into subspaces based on how long ago the information, such as a cue-similarity map or fixation location, was stored. Each subspace is dynamically updated as the task progresses, maintaining information according to its temporal distance from the current time step (Fig. 7B; right).

**Figure 7.**
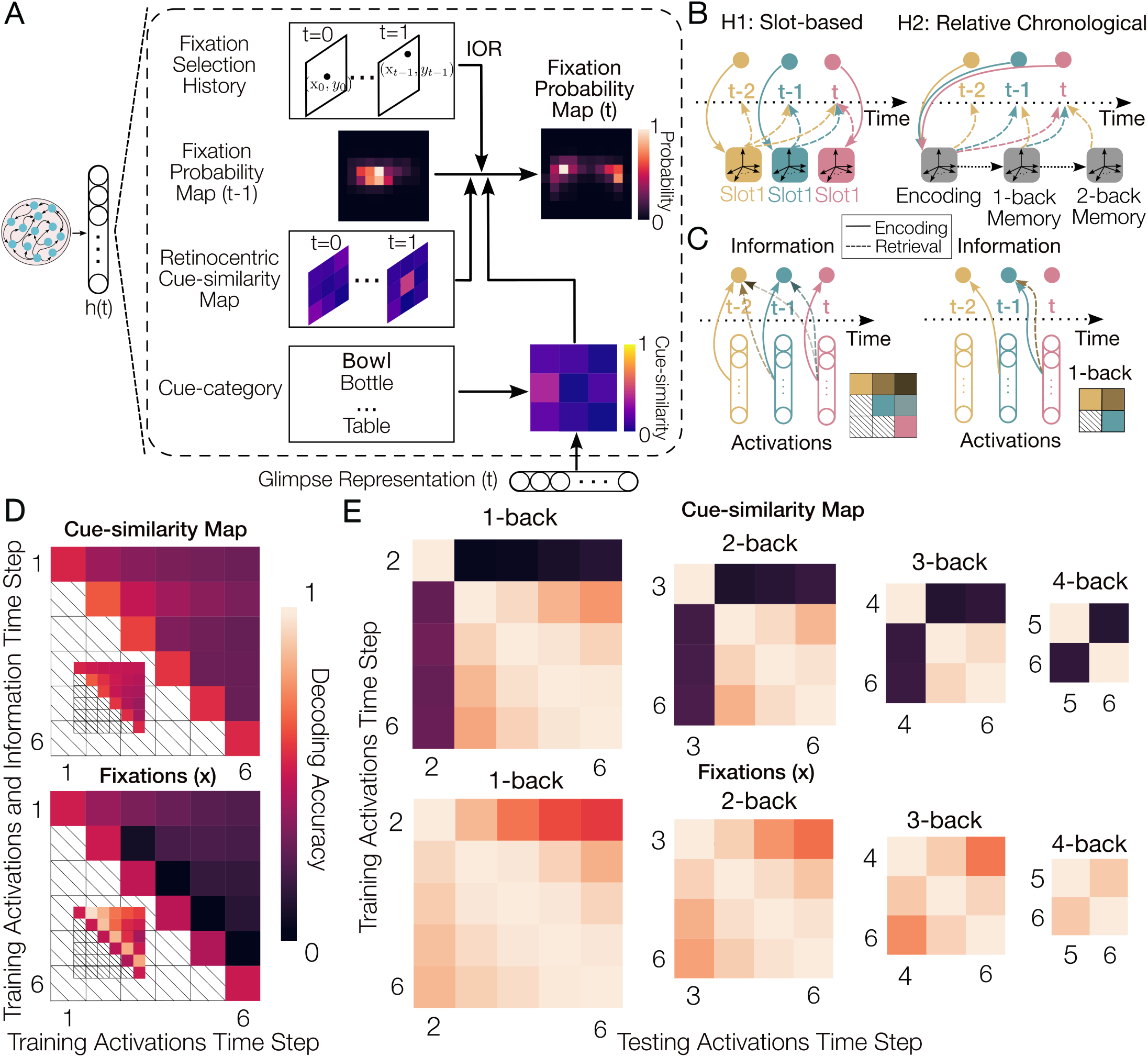
Neural mechanisms of maintenance and retrieval of memory during visual search. A. Schematic of latent representations guiding fixation selection in GVSM. Retinocentric cue-similarity maps are recomputed at each step from stored cue-category information and incoming glimpses. The model encodes both current and past cue-similarity maps, along with fixation selection history. These elements are integrated to update an allocentric fixation probability map that guides future fixations. Fixation selection history contributes to this update through an inhibition of return (IOR) mechanism. **B. Schematic of two candidate memory encoding mechanisms.** H1: Slot-based memory: Each item is stored in a distinct, temporally stable subspace of the latent space. H2: Chronologically indexed memory: Items are stored in subspaces indexed by relative temporal distance, with memory contents dynamically updated as time progresses. **C. Schematic of decoding procedures used to evaluate memory storage mechanisms.** Left: To test the slot-based hypothesis, linear decoders are trained to decode information (e.g., retinocentric cue-similarity maps or fixation coordinates) from the hidden state at the time the information becomes available (solid arrows). These slot decoders are then tested at later time steps to assess whether the same encoding subspace is preserved over time (dashed arrows). Decoding accuracy is visualized in a heatmap, where each cell (i, j) reflects the performance of a decoder trained at time step i (encoding time) and tested on activations from time step j. High off-diagonal accuracy (i ̸= j) would indicate that the information remains encoded in a stable, time-invariant subspace. Right: To test the chronologically indexed hypothesis, linear decoders are trained to decode information stored at a fixed temporal distance in the past (e.g., 1-back), regardless of its absolute encoding time (solid arrows). These decoders are then tested at other time steps to assess whether the same relative-time subspace (e.g., 1-back) is preserved across the trial (dashed arrows). In the decoding matrix, the diagonal elements (i, i) represent the decoder trained and tested on decoding the same n-back information (e.g., 1-back), while the off-diagonal elements (i, j) show generalization of the n-back decoder across time—indicating whether the subspace is consistently indexed by relative temporal distance. **D. Lack of slot-based memory subspaces.** The main heatmaps show decoding accuracy for cue-similarity map representations (top) and fixation *x*-coordinates (bottom), using decoders trained at each encoding time step and tested across subsequent time steps (as schematized in Fig. 7C, left). Diagonal elements reflect encoding accuracy, while off-diagonal elements quantify retrieval accuracy using the original (slot-based) subspace. In both cases, decoding accuracy drops along each row, indicating that the subspace used to encode information is not preserved over time. The inset shows baseline memory decoding: each (i,j) element quantifies how well the information from time step *i* is still present (linearly decodable) at time step *j*, irrespective of subspace alignment. Notably, the drop in slot-based retrieval is greater than the loss of memory content, confirming that the encoded information is retained but its subspace changes—ruling out the presence of slot-based memory subspaces for both cue-similarity and fixation history. **E. Memories are encoded in relative chronological subspaces after the first step.** We tested whether memory representations are organized by their relative temporal distance from the current time step (as schematized in C, right). Four heatmaps show decoding accuracy for 1-back to 4-back information. Each column is normalized by its diagonal element, which reflects the maximum decodable amount of that memory content at a given time step (i.e., the ceiling for that column). After the first time step, decoders generalize well across time for both cue-similarity maps (top) and fixation *x* coordinates (bottom), indicating that these memories are stored in relative chronological subspaces.

To test these hypotheses, we performed decoding analyses using linear regression models on both the memory of the cuesimilarity, and fixation selection history. We focused on the *x*-coordinates decoding only, as the *y*-coordinates exhibited strong correlation across time, resulting in a shared encoding subspace across all time steps (Fig. 3). This prevented a meaningful analysis of *y*-coordinate memory mechanisms. To test H1, we fitted linear regression models to decode memory content from the unit activations at the time step in which the information became available (i.e., the encoding time). We then assessed the generalization of these decoders to later time steps (Fig. 7C; left). High generalization accuracy would suggest that the information remains stably encoded in the same subspace, consistent with a slot-based mechanism. Conversely, low generalization would indicate that the encoding subspace is not preserved over time. To test H2, we fitted linear regression models to decode n-steps-back information from unit activations at each time step and evaluated their generalization to decoding the same n-steps-back information at other time points (Fig. 7C; right). High generalization accuracy in this analysis would support the presence of subspaces organized by relative temporal distance, consistent with chronologically indexed memory subspaces.

We found that none of the cue-similarity map or fixation memories are stored in slot-based subspaces, as the encoding subspace did not encode the same information at later time steps, i.e. drop from diagonal to off-diagonal elements in each row (Fig. 7D). We ruled out the possibility of this drop being due to weaker memory representations compared to encoding representations by comparing with the baseline of memory decoding, which is quantifying how much past information is preserved within the latent space at later time steps (Fig. 7D; insets, and Fig. 3 E-F). We found that after the first time step, the memory is stored in the relative chronological memory subspaces indicated by the fact that n-back decoders trained on one time step generalize well to the other time steps (Fig. 7E). However, the memory of the first step is not encoded in a relative chronological subspace. After encoding, at later time steps, it is transformed via rotation into a new subspace and remains stored there.

Overall, our results revealed distinct memory storage mechanisms for the first time step and the later ones, for both the cue-similarity map and fixation memory. The first time step information was transformed into a new subspace and stored there consistently throughout the trial, whereas the memory of the later time steps was stored in relative chronological subspaces.

## Discussion

Much of the recent progress in brain-predictive modeling has focused on early visual areas, where feedforward neural networks trained on static object recognition tasks have successfully predicted neural responses across the ventral stream. While large language models have extended this approach to higher order cortical regions involved in language processing, few models have tackled the dynamic, interactive computations required for active vision. Visual search exemplifies such behavior: it is a goal-driven task that engages multiple cognitive functions, including selective attention, working memory, and sequential decision-making, and has long served as a central paradigm in the study of higher-order visual cognition. Yet traditional artificial neural network models of vision typically rely on uniform retinal sampling and static images, offering no mechanism for interaction with the environment. As a result, predictive modeling of downstream fronto-parietal areas such as FEF, VPA, and LIP, regions critical for memory and attention-guided eye movements, has remained largely unexplored. To address this gap, we developed a task-optimized model of visual search that operates directly on natural scenes and learns to guide sequential foveated sampling. This model, GVSM, replicates both behavioral patterns and neural response properties observed in these higher-order cortical areas. By extending predictive modeling beyond the sensory cortex into the realm of active vision and cognitive control, GVSM offers a framework for studying the emergent mechanisms that underlie complex natural behavior.

To this end, we developed a neural network model of visual search, called GVSM, addressing key limitations of prior models, namely, lack of a dynamic internal state for integrating information across fixations, reliance on symbolic rules rather than learned latent representations, and unrealistic assumptions about uniform visual acuity or biologically implausible retinal transformations. GVSM features a biologically inspired architecture that searches for targets directly from pixel inputs, making it both image-computable and neurally plausible. It incorporates eccentricity-dependent visual acuity, maintains a dynamic internal representation to integrate information across fixations, and generates artificial neural activations that are testable against brain data. Trained solely on the visual search task objective, GVSM exhibits strong generalization to unseen stimuli and captures both high-level and fine-grained aspects of human search behavior, producing human-like search scanpaths. Furthermore, the model adopts human-like search strategies and encodes information using brain-like neural representations. We found that its fixations are predicted by cue similarity and fixation history in ways that closely mirror human behavior. GVSM addresses several limitations of previous visual search models. It integrates eccentricity-dependent visual acuity, maintains a dynamic internal state across fixations, and learns flexible, latent representations for fixation selection directly from pixel inputs. Trained solely to maximize task performance, GVSM generalizes well to unseen scenes and categories, produces human-like scanpaths, and recovers core signatures of visual search observed in human behavior and primate physiology. Importantly, because GVSM is not explicitly hand-engineered with mechanisms such as inhibition of return, it naturally integrates fixation history into its fixation selection process, resulting in fixation dynamics that are more aligned with those observed in humans.

### Human-like scanpaths and emergent biases

A hallmark of visual search behavior is the spatiotemporal sequence of fixations, scanpaths, that observers generate while searching for a target. GVSM closely replicates these scanpaths on a trial-by-trial basis, producing fixation sequences that mirror human behavior in both their structure and variability. Beyond this, the model captures a wide range of human-like saccadic statistics, including saccade size distributions, directional preferences, and the dynamics of fixation transitions, despite lacking any explicit oculomotor constraints. Notably, GVSM exhibits a strong horizontal bias in its fixations, a robust feature of human search behavior, despite not being constrained by biomechanical or motor factors. This behavioral horizontal bias was also reflected in the latent representations, indicated by higher decodability of the *y* coordinates of upcoming fixations than the *x* coordinates. Interestingly, this asymmetry is only present at the selection stage, and after a fixation is made, memories of both its *x* and *y* coordinates are equally retrievable from the latent space at later time steps.

This emergent horizontal bias supports the hypothesis that directional preferences arise from ecological regularities in natural scenes and the statistics of visual behavior, rather than from physical limitations Tatler and Vincent (2009). Everyday behaviors such as reading and exploration, which unfold predominantly along the horizontal axis, may have shaped both the human visual system and the visual input used to train our model, contributing to the emergence of horizontal bias. Consistent with this interpretation, the model also captures other human-like saccadic properties such as saccade size, direction distributions, changes in saccade direction, and fixation eccentricities, despite lacking an oculomotor system.

### Neural correlates of working memory and attention

GVSM also recapitulates neural correlates of memory and attention that have been extensively documented in fronto-parietal areas. Units in its Information Integrator module show persistent category-selective responses during the cue delay period, similar to sustained activity observed in VPA. Feature and spatial selection effects likewise emerge, with unit responses modulated by the presence of a target in the RF and the direction of the upcoming saccade, echoing findings from FEF and LIP. These results suggest that the model implicitly learns to prioritize targets in a way that mimics attention-related modulation in primate fronto-parietal circuits.

A particularly striking result is the spontaneous emergence of a distributed retinocentric cue-similarity map in GVSM’s latent space. Even though the model is trained only to perform the task, it encodes cue-similarity values across the visual field in a manner consistent with neurophysiological findings. This map guides fixation selection and is maintained in memory across time, consistent with theories positing distributed topographic encoding of visual priorities in areas such as FEF, VPA, and LIP.

### Structured encoding and dynamic transformation of visual memory

Beyond recovering population-level signa-tures, GVSM also offers insights into the geometry of the neural code underlying visual search. Cue-similarity representations were organized topographically: nearby locations in the visual field were encoded in more similar subspaces, and generalization performance declined with increasing spatial distance.

Interestingly, the encoding subspace underwent a sharp transformation after the first glimpse of the scene. Cross-temporal generalization and Procrustes analysis revealed that this shift could be captured by a rotation of the encoding axes, after which the representation stabilized. This transition suggests that early visual input is encoded in a distinct subspace that is rapidly reformatted to support subsequent search, a form of representational processing not explicitly imposed by the model’s design.

Ablation studies further revealed that much of the model’s search trajectory is planned following the initial scene glimpse. When visual input was occluded after the first step, GVSM still performed above chance, indicating that early observations suffice to guide several fixations. However, performance declined as the number of required fixations increased, consistent with a shift from preplanned to reactive control. These results align with behavioral and physiological evidence suggesting that visual search integrates both memory-based planning and online sensory input.

### Toward neurally grounded models of cognitive function

GVSM offers a testable, neurally grounded account of visual search that bridges behavior and brain-like representations. The model generates several predictions: (1) cue-similarity is encoded in a continuous topographic format, with subspace alignment decreasing with spatial distance; (2) this representation rotates over time, transitioning from an initial subspace to a stable format while preserving internal geometry; (3) cue category information is maintained in a stable subspace across the trial; and (4) memory of past fixations and cue-similarity maps is stored in relative chronologically indexed subspaces after the first time step while the memory of the first time step is preserved in a slot transformed from its encoding subspace. These findings provide concrete hypotheses for future neurophysiological studies.

### Extending predictive modeling beyond the sensory cortex

Much of the success of artificial neural network models in neuroscience has centered on sensory systems, where deep networks trained for object recognition Yamins et al. (2014); Ratan Murty et al. (2021); Bashivan et al. (2019); Cadieu et al. (2014); Khaligh-Razavi and Kriegeskorte (2014), auditory classification Kell et al. (2018), or odor identification Wang et al. (2021); Taleb et al. (2024) have reproduced cortical response patterns in ventral visual, auditory, and olfactory areas. More recently, large language models have shown promise in predicting activation patterns across the cortical language network, including Broca’s area, the posterior superior temporal gyrus, and angular gyrus Caucheteux and King (2022); Schrimpf et al. (2021). Yet, these models have rarely been extended to predict neural responses in higher-order cortical regions such as those in the prefrontal or parietal cortex, areas involved in flexible, task-dependent control, likely due to the broader behavioral repertoire and complexity of the computations performed there.

GVSM takes a first step toward this goal. By aligning task optimization with the dynamics of working memory, attention, and decision-making, it offers a computational model of fronto-parietal function grounded in a naturalistic behavior. The model’s architecture generalizes beyond symbolic rules or hard-coded dynamics, making it extensible to other cognitive domains. As large-scale neural recordings during naturalistic visual search become available, our model can be directly tested for its predictive power in these areas. Furthermore, the general architecture of our model makes it a promising foundation for extension to other cognitively demanding tasks, offering a path toward unified models of high-level cognitive function that can predict neural responses across the prefrontal and parietal cortices, moving beyond the sensory and language domains into the realm of attention, memory, and decision-making. As predictive models continue to advance across domains, our work highlights the value of using goal-driven architectures not merely as engineering tools, but as scientific instruments to probe the computational principles of perception and action in the brain.

## Methods

The model was implemented and trained in Python (v3.12; Python Software Foundation, https://www.python.org/) using the PyTorch library (v1.14; https://www.pytorch.org/) on a GPU-accelerated machine equipped with 1–4 NVIDIA RTX A5000 graphics cards. All analyses of human and model data were conducted in Python v.3.12, using the numpy v.1.26.4 and/or scikit-learn v.1.5.2 packages.

### Dataset

#### Training Dataset: Large-Scale Natural Scene Visual Search Dataset Generation

To train GVSM, we constructed a large-scale natural scene visual search dataset based on the Places365 image dataset, which contains over 10 million images of diverse real-world environments Zhou et al. (2017). Each sample in the dataset consists of an arbitrary instance of a cue category (e.g. car), a visual scene that contains at least one instance of the cue category, and a ground truth mask that specifies the position of the cue category in the given scene. To generate the ground truth target masks, we processed each image with a state-of-the-art object detection model (Mask R-CNN) He et al. (2017) and extracted object masks for all detected instances of 80 cue categories (e.g., car, table, kite, person). The resulting cue bank comprised an average of 62,138±257,067.33 exemplars per category, with ”person” having the largest set (2,261,195 exemplars) and ”hairdryer” the smallest (230 exemplars) (Fig. 1B). To simplify the training and avoid multiple simultaneous targets, we merged all object masks of the same category within an image into a single composite mask. Consequently, any region containing an instance of the target cue category was considered part of the target area.

Each (image, cue category) pair served as a distinct trial. The order of presenting the trials was randomized on each pass through the dataset. Task frames were generated on the fly: the cue frame was created by placing a randomly selected exemplar from the cue bank at the center of a black 640 × 640 pixels canvas, while the search image was resized such that its longer side matched 640 pixels, with the shorter side padded symmetrically to yield a 640 × 640 pixels search frame. Ground-truth target maps were generated by overlaying a 10×10 grid on the cue object mask, thereby discretizing the visual field into 100 spatial bins and reducing the model’s action space from pixel-level to grid-level selection. During the initial training phase (supervised training), we used strict ground-truth maps, marking a grid location as a target only if it was fully covered by the object mask. In a subsequent training phase, we adopted a more lenient labeling approach, similar to other visual search models, where any grid cell with partial overlap with the target area was marked as a hit. The final dataset comprised 6,754,405 total trials, drawn from 4,472,151 unique images and 80 cue categories.

#### Testing Dataset: COCO-Search18

To evaluate the behavioral alignment between GVSM and human visual search behavior, we used the COCO-Search18 dataset Chen et al. (2021). This large-scale benchmark comprises over 6,000 natural images (target-present only) from the MS COCO dataset, each annotated with eye movement trajectories recorded during visual search. The dataset includes data from 10 human participants who were instructed to search for one of 18 object categories (e.g., microwave, stop sign, backpack) per trial. Each trial contains the sequence of fixations, their durations, and the identity of the search target, offering a detailed account of goal-directed attention and saccadic decision-making in complex real-world scenes.

To generate corresponding search trials for our model, we adapted the same pipeline used for training data generation. Instead of text queries (as used in the human experiments), we constructed cue frames by presenting an exemplar image of the target category at the center of a black 640×640 pixels background. We also derived lenient ground-truth target maps for the model using the object bounding boxes provided in the dataset, assigning a grid location as a target if it overlapped with the ground-truth target area. Our evaluation focused exclusively on the target-present subset, and we assessed both spatial and temporal alignment between the model’s predicted fixation sequences and human scanpaths.

### GVSM

#### Retinal transformation

To approximate the progressive loss of visual acuity in the periphery caused by the retinal transformation, we implemented a multi-resolution cropping module adopted from prior work Mnih et al. (2014). For each 640 × 640 task frame, the module extracts three square crops centered at the model’s current fixation location with sizes of 128, 256, and 512 pixels. All three crops are then resized to 128 × 128 pixels. The original 128 × 128 pixels crop serves as the high-resolution foveal input, while the resized 256 × 256 pixels and 512 × 512 pixels crops provide increasingly blurred peripheral representations. This setup mimics the eccentricity-dependent degradation of visual detail in the human retina.

#### Model architecture

The model consists of four main components: a visual processing module, a glimpse network combining the visual observation with the current fixation location, an information integration module, and a policy network. In the visual processing module, the retinally transformed images are processed by an EfficientNet-B0 CNN Koonce (2021), which simulates the visual processing along the ventral visual pathway Schrimpf et al. (2018). From each input image, we extract the final convolutional feature maps before the last fully connected layer, yielding a 1280 × 4 × 4 representation of the scene. This visual representation is then combined with the current fixation location through a feedforward network denoted as the glimpse network. Fixation coordinates (*x*,*y*) are defined in a normalized allocentric reference frame, with the image center as the origin. The x-axis increases rightward and the y-axis upward, with both coordinates scaled to lie in the range [-1,1]. In the glimpse network, the fixation location and the CNN-derived visual features are separately projected to half the embedding dimension (512) using linear layers followed by ReLU nonlinearities. These two projected vectors are then independently transformed by additional linear layers to the full embedding dimension (1024), summed, and passed through another ReLU, yielding the glimpse representation.

The glimpse representation is processed by the Information Integrator module, a 1024-unit LSTM or an 8-layer causal transformer with 8 attention heads and an embedding dimension of 1024. The Information Integrator’s latent space, LSTM hidden state or the final output token of the transformer, is passed on to the policy network which is a multi-layer perceptron (MLP). This network, composed of 3 fully connected layers with *tanh* nonlinearity, outputs a 100-dimensional probability distribution over the 10 × 10 spatial grid. The next fixation location is sampled from this distribution. This new fixation location is then passed to the retinal transformation module to generate a foveated input for the next time step. The foveated input re-enters the CNN, initiating another pass through the model. This loop continues until the model either fixates on the target area or reaches the maximum number of time steps (10). The first two steps are fixed: the model first fixates on the cue frame, then at the center of the search image. Thus, the model is allowed to make up to 8 fixation decisions per trial.

### Model Training

We first pretrained the CNN to perform object recognition on images from the ImageNet dataset, each subjected to a simulated retinal transformation centered at randomly sampled fixation locations. The task was to classify each image into one of 1,000 categories. Training hyperparameters followed standard practice in the CNN literature, and in particular, the specifications introduced in the original EfficientNet-B0 paper Koonce (2021) adjusted to the available resources (1 GPU: Nvidia V100SXM2) (maximum learning rate = 0.36, weight decay = 10*^−^*^5^, learning rate scheduler = 5 epochs warmup followed by waterfall scheduling, multiplying the learning rate by 0.97 every 2.4 epochs, optimizer = RMSProp Tieleman (2012), batch size = 512). We then adopted a two-phase training paradigm inspired by prior work on saccade-augmented visual categorization Elsayed et al. (2019). This approach mirrors the two-stage training framework used in recent foundation models, such as large language models, in which an initial self-supervised learning phase (here, supervised) is followed by an RL stage Christiano et al. (2017); OpenAI Achiam et al. (2023); Touvron et al. (2023); Baker et al. (2022).

#### Phase 1: Supervised training

In the first phase, we optimized the parameters of the LSTM/transformer (with CNN parameters kept frozen) to predict the target area given a random sequence of fixations over the search image. A linear readout layer was applied to the LSTM hidden state/final output token of the transformer to produce a 100-dimensional vector, representing predicted probabilities over a 10×10 grid. This output was trained using binary cross-entropy loss against the strict ground-truth target maps. To address the class imbalance, since most grid locations in a trial do not contain the target, we introduced per-sample, per-class weighting to balance the contribution of positive (on-target) and negative (off-target) locations. The weighted binary cross-entropy loss is defined as:

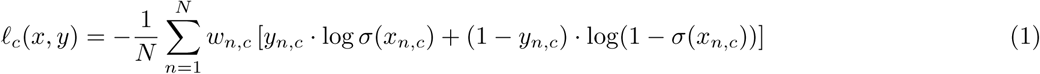

where *σ*(.) is the sigmoid function, *N* is the batch size, *C* = 100 is the number of classes (grid locations), *x* ∈ *R^N×C^* are the model logits, *y* ∈ {0, 1}*, N* × *C* are the binary labels (1 if on the target, 0 otherwise), *w_n,c_*is the weight for sample *n* and class *c* defined as:

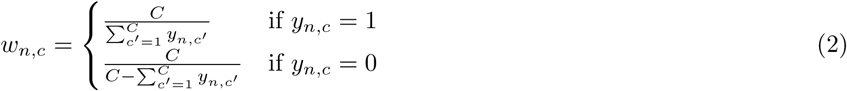

Here, 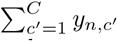 is the number of positive labels (on-target) for sample n. This formulation ensures comparable loss magnitude across images with large differences in target sizes. for each trial, the total loss contribution from on-target and off-target locations is balanced, regardless of their relative frequency.

#### Phase 2: reinforcement learning

In the second phase, we trained the MLP policy network to sequentially choose fixation locations based on the LSTM hidden state/transformer’s final output token. During this stage, the parameters of the CNN and information integrator (LSTM/transformer) were frozen. The MLP received the 1024-dimensional LSTM hidden state/transformer’s latest output token and produced a 100-dimensional probability distribution over the grid locations. The fixations were sampled from this distribution over grid locations using softmax sampling. We used the Proximal Policy Optimization (PPO) algorithm Schulman et al. (2017) to optimize the policy network. A reward of +10 was assigned if the selected fixation landed on the target area (as defined by the lenient target map), and a penalty of -1 was given for each step taken without finding the target. This reward structure encouraged the model to locate the target as efficiently as possible. To maximize training efficiency and avoid uninformative episodes, we curated a subset of the trials from Phase 1 for RL. We excluded trials in which the target was excessively large or small, as oversized targets can inflate cumulative rewards without improving performance in realistic conditions. Target size thresholds were matched to the minimum and maximum target sizes observed in COCO-Search18. We also limited cue categories to the 18 used in COCO-Search18. Despite being trained on this curated subset, the RL agent successfully generalized to the full dataset used in Phase 1, owing to its exposure to the most informative and representative trials during training.

### Behavioral and Latent Space Analysis

#### Baselines

To ensure a fair comparison between our model and baseline models, we closely matched the target-found criteria and other evaluation conditions to those used in GVSM. First, we aligned the fixation box size across models. The fixation box refers to the area around the fixation point that is considered “seen.” If this area overlaps with the target, the search is deemed successful; otherwise, the area is inhibited, and its probability of being fixated again is set to zero. While our model does not implement explicit inhibition of return, its action space is restricted to a 10 × 10 grid, making each 64 × 64-pixel location analogous to fixation boxes in IVSN and eccNET, and to grid cells in IRL. To standardize evaluation, we matched the fixation-box-to-image ratio across all baselines to that of our model (64/640 = 0.1). Using the original fixation sizes of the baselines did not meaningfully alter results. We used the publicly available implementations of the baseline models and the fixation-by-fixation scanpath prediction method provided by Travi et al. (2022). Note that IRL received cue categories in textual form, while IVSN, eccNET, and our model used visual exemplars of the cue. To further ensure consistency, we mapped all scanpaths, human and model, to a common reference frame: a 640 × 640 pixel image space discretized into a 10 × 10 grid. We reduced consecutive fixations falling within the same grid cell to a single fixation at that location. The same target-found criterion was applied: if the fixated grid location overlapped with the target region, the target was considered found and the trial ended.

#### Fixation-by-fixation evaluation of search scanpath predictions

To evaluate how well the models predicted human visual search behavior at a fine-grained level, we used a fixation-by-fixation scanpath prediction method Kümmerer and Bethge (2021). This approach represents the state-of-the-art in scanpath comparison and addresses several critical limitations of traditional sequence-based metrics, which typically treat scanpaths as holistic sequences and rely on distance-based similarity measures. In contrast, the fixation-by-fixation approach (1) better reflects the decision-making process underlying eye movements by treating each fixation as a choice conditioned on previous ones, (2) aligns with standard evaluation methods used in saliency map prediction, (3) enables detailed behavioral alignment analyses by scoring individual fixations rather than entire scanpaths, and (4) is free of hyperparameters.

#### Comparing distributions of fixation directions

To compare the distribution of fixation directions between models and humans, we computed the Jensen–Shannon (JS) divergence between their respective direction histograms Lin (2002). Given two normalized distributions P (model) and Q (human), the JS divergence is defined as:

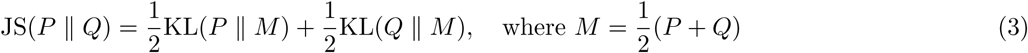

where the Kullback–Leibler (KL) divergence is given by Kullback and Leibler (1951):

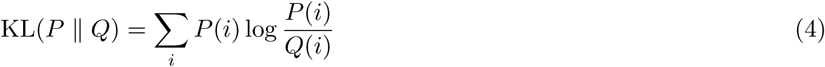

Unlike KL divergence, which is asymmetric and can be undefined when Q(i)=0, JS divergence is symmetric, smooth, and always finite. These properties make it a robust and interpretable measure of dissimilarity between distributions. A lower JS divergence indicates that the model’s fixation direction distribution more closely resembles that of humans.

#### Computing the cue-similarity maps

To assess the similarity between the cue object and various locations on the search image, we convolve the visual representation of the cue frame (i.e., the final feature maps of the CNN before the global pooling and the classifier fully connected head) with the image Zhang et al. (2018). Map is computed in a retinocentric framework, i.e. the images are retinally transformed. They, 3-channel 128×128 (grayscale) input, are processed by the EfficientNet-B0 CNN. The CNN is the same as the one used as the vision module in the model, pretrained on retinal-transformed ImageNet to do object recognition. The resulting cue-similarity maps are 3×3.

#### Latent decoding analysis

To examine the representational content of the model’s latent state, we conducted linear decoding analyses using ridge regression. We fitted linear decoders to predict various target variables, including cue-similarity maps and (*x*,*y*) coordinates of visited locations from the model’s latent activations at each time step. Specifically, for each model instance, a separate decoder was trained for each spatial location (i.e., each cell in a 10×10 grid over the image), using the output token of the Information Integrator module as input features. Fitting was performed with 5-fold cross-validation, and decoding accuracy was quantified as the Spearman correlation between predicted and ground truth values on held-out trials. To evaluate subspace generalization, such as from cue-similarity maps to fixation probability maps, across time steps, or across spatial locations, we tested trained decoders on mismatched targets and compared their performance to within-target decoding baselines.

#### Orthogonal Procrustes analysis

To examine the geometrical structure of neural representations in the model’s latent space, we used orthogonal Procrustes analysis to quantify how well one set of decoder weight vectors could be aligned with another via rotation. Each set of weights, corresponding to either cue-similarity maps or fixation probability maps, comprised 100 vectors (one per 10 × 10 grid cell), trained to predict the value of the respective spatial map at each location based on the latent activation at a given time step. We used this framework to assess whether the representation of cue-similarity maps rotated across time, i.e., whether the encoding subspace at one time point could be aligned to that of another via a rigid rotation. To do so, we first centered and normalized the source and target sets of weight vectors. We then applied orthogonal Procrustes analysis to compute the optimal rotation matrix and global scaling factor, aligning the source to the target. The transformed source weights were then inverse-transformed to recover their original scale, and alignment quality was quantified by evaluating the decoding performance of the reconstructed decoders on the target prediction task. This approach allowed us to identify the geomteric transformation of the cue-similarity representations across time.

## Acknowledgments

This research was supported by the Healthy-Brains-Healthy-Lives startup supplement grant, the NSERC Discovery grant RGPIN-2021-03035, and CIHR Project Grant PJT-191957. P.B. was supported by FRQ-S Research Scholars Junior 1 grant 310924, and the William Dawson Scholar award. M.P. was supported by the UNIQUE Masters and PhD Fellowships, and the Stichting Formation Award. All analyses were executed using resources provided by the Digital Research Alliance of Canada (Compute Canada) and funding from Canada Foundation for Innovation project number 42730.

